# Environment and host infection history jointly predict disease risk in a multi-pathogen system

**DOI:** 10.64898/2026.04.13.718236

**Authors:** Carly B. Scott, Susan Cleary, Rita L. Grunberg, Fletcher W. Halliday, Brooklynn N. Joyner, Kayleigh R. O’Keeffe, Isabelle Stiver, Charles E. Mitchell

## Abstract

Forecasting disease epidemics may require considering both abiotic and biotic conditions. Abiotic conditions can influence pathogen dispersal, survival, and infection, while prior infection by one pathogen species can alter host susceptibility to subsequent pathogens, creating historical contingencies. Yet the relative importance of environmental conditions, within-host pathogen interactions, and their interplay in predicting seasonal disease dynamics remains underexplored. We analyzed approximately 30,000 plant-level observations of three foliar fungal diseases on the grass species tall fescue in a North Carolina old field over seven years using three complementary modeling approaches: random forest, survival analysis, and Bayesian hierarchical models. Temporally explicit random forest models predicted future disease with over 80% accuracy for two of the three diseases and revealed environmental thresholds for disease prediction in abiotic variables including temperature and humidity. Survival analyses showed that pathogen-pathogen interactions rivaled environmental conditions in predicting risk of crown rust, which was strongly facilitated by prior anthracnose infection. Across all pathogens, Bayesian hierarchical models recovered evidence that facilitative interactions between coinfecting pathogens were modulated by effects of environmental conditions, particularly temperature. Together, these complementary approaches show that a host’s current disease state can matter as much as its environment in predicting disease, identify key environmental factors associated with disease risk, and show that modulation of environmental drivers by pathogen coinfections can improve predictions of seasonal disease epidemics. Other disease forecasts may be improved by accounting not just for environmental drivers, but also how those drivers propagate through pathogen species interactions.

**Open Research Statement:** All data and code used to generate the results contained in this paper are available on GitHub (https://github.com/cb-scott/WidenerLongTermAnalysis) and archived on Zenodo at 10.5281/zenodo.22285642.

## Introduction

Pathogen transmission success often depends on both abiotic environmental factors and biotic context. Abiotic conditions, such as temperature, moisture, and wind, can each shape a pathogen’s ability to survive, disperse, and infect a host (Altizer et al., 2006). Epidemics are often linked to seasonal environmental cycles of these variables (Penczykowski et al., 2015), acting either through the pathogen’s own physiology, for example infectivity, or through host physiology, such as susceptibility to infection (Chen et al., 2024; Garrett et al., 2021).

Beyond these abiotic filters, hosts present a biotic landscape that pathogens must navigate. Colonizing pathogens encounter an established microbial community, potentially including other pathogens that have already infected the host. These interactions can be facilitative or antagonistic and are often modulated by host immunity and the pathogen’s life history strategy. Earlier infection by one pathogen may "prime" defenses against subsequent infection by other pathogens (Goellner & Conrath, 2008) or modify the amount of nutrient available in host tissue (Wale et al., 2017), reducing the success of later arrivals. In contrast, pathogens that trigger different host immune responses may facilitate each other through immune crosstalk (Pieterse et al., 2012).

These within-host pathogen-pathogen interactions have the potential to alter disease outcomes and create historical contingencies that may scale up to the level of the host population (Fukami, 2015; Susi et al., 2015). The outcome may depend not only on coinfection status, but also on the order in which pathogens arrive (Halliday et al., 2017). For example, ignoring arrival order of bacterial and fungal diseases in a model of zooplankton disease qualitatively failed to capture multi-pathogen epidemic dynamics altogether (Clay et al., 2020), highlighting that earlier-arriving pathogens can fundamentally reshape the infection landscape for those that follow.

The abiotic environment may further modulate the strength and direction of pathogen-pathogen interactions (Hanke et al., 2015; Jiang et al., 2011). For example, seasonal temperature variation determined whether an endophytic biocontrol bacterium suppressed or failed to suppress the bacterial pathogen, *Ralstonia solanacearum*, over the tomato growing season (Wei et al., 2017). Similarly, across systems, the relative importance of abiotic and biotic context may shift as conditions change through the season. However, the relative contributions of environmental variation and pathogen-pathogen interactions to seasonal disease epidemics, as well as the importance of interactions between abiotic and biotic variables, remain poorly resolved (Alexander et al., 2016; Halliday et al., 2021; Suh et al., 2024).

Resolving the roles of abiotic conditions and biotic interactions while both are varying over time may benefit from the integration of multiple, complementary modeling approaches. Random forest algorithms accommodate collinearity and non-linear relationships among predictors and can reveal thresholds at which abiotic drivers shift disease outcomes (Alves et al., 2025; Shah et al., 2023). However, these algorithms prioritize predictive accuracy, so mechanistic insight may require other approaches. In disease systems, survival analyses offer a natural complement, accommodating repeated sampling of individuals through time and handling censored time-to-event data (Scherm & Ojiambo, 2004). Bayesian hierarchical models offer a similar advantage, directly modeling complex experimental designs (Farnsworth et al., 2006; Gibson et al., 2004). Combined, these approaches may balance predictive power with mechanistic insight grounded in the full experimental design.

Here, we analyze seven years of data on three plant foliar fungal diseases, crown rust, brown patch, and anthracnose, on the grass species tall fescue in an old field. Previous studies conducted over a one- to three-year time frame show that interactions between these pathogens in tall fescue can alter disease progression (Green et al., 2024; O’Keeffe et al., 2021) and influence host susceptibility to later-arriving diseases (Halliday et al., 2017). Building on these short-term experiments, we leverage multi-year variation to quantify how the influence of pathogen–pathogen interactions compare with abiotic context in shaping seasonal epidemics.

To accomplish this, we integrate three complementary modeling approaches to examine how abiotic conditions and host infection history jointly shape disease risk. First, we use random forest models to identify key environmental predictors and potential thresholds in disease risk from a complex set of correlated abiotic variables. Second, we apply survival analyses to a subset of plants in one year that were monitored longitudinally to evaluate how abiotic conditions and prior infection by other pathogens influence subsequent disease risk at the plant level. Finally, we use Bayesian hierarchical models to leverage the full seven-year dataset and account for repeated sampling within plots, allowing us to test how the effects of pathogen coinfection vary depending on environmental conditions at the plot level. Together, these approaches reveal that host infection history can be as important as abiotic conditions in predicting disease risk, and that the consequences of prior infection depend on environmental context. Our findings highlight how interactions between pathogen species and their abiotic environment jointly shape disease dynamics.

## Methods

### Study System

Tall fescue (*Lolium arundinaceum*) is a widespread cool-season grass that is grown for turf, fodder and forage, and is an ecologically dominant species in many unmanaged plant communities across much of the eastern United States (Natural Resources Conservation Service, 2001). We studied diseases of tall fescue in Widener Field of Duke Forest Teaching and Research Laboratory, in the Piedmont region of North Carolina. Widener Field was used to grow row crops until 1996; since then, it has been maintained as an herbaceous-dominated old field by annual mowing.

We focused on three foliar fungal diseases which predominantly infected tall fescue at our field site: anthracnose, brown patch, and crown rust. Anthracnose at our field site is caused chiefly, but not exclusively, by *Colletotrichum cereale*, which spreads primarily through rain-splash dispersal of asexual conidia (Crouch & Beirn, 2009). Brown patch is caused by *Rhizoctonia solani*, which persists in the soil, and disseminates through hyphal fragments and growth in soil and water films as well as sclerotia (Ogoshi, 1987). Crown rust is caused by *Puccinia coronata* and is dispersed via wind-borne urediniospores. Together, these diseases affect foliar health and performance of tall fescue across diverse environmental conditions (Butler & Kerns, 2019a, 2019b, 2019c). All three diseases are visually distinct from each other in the field and were identified based on these characteristic symptoms (Figure 1).

**Figure 1.**
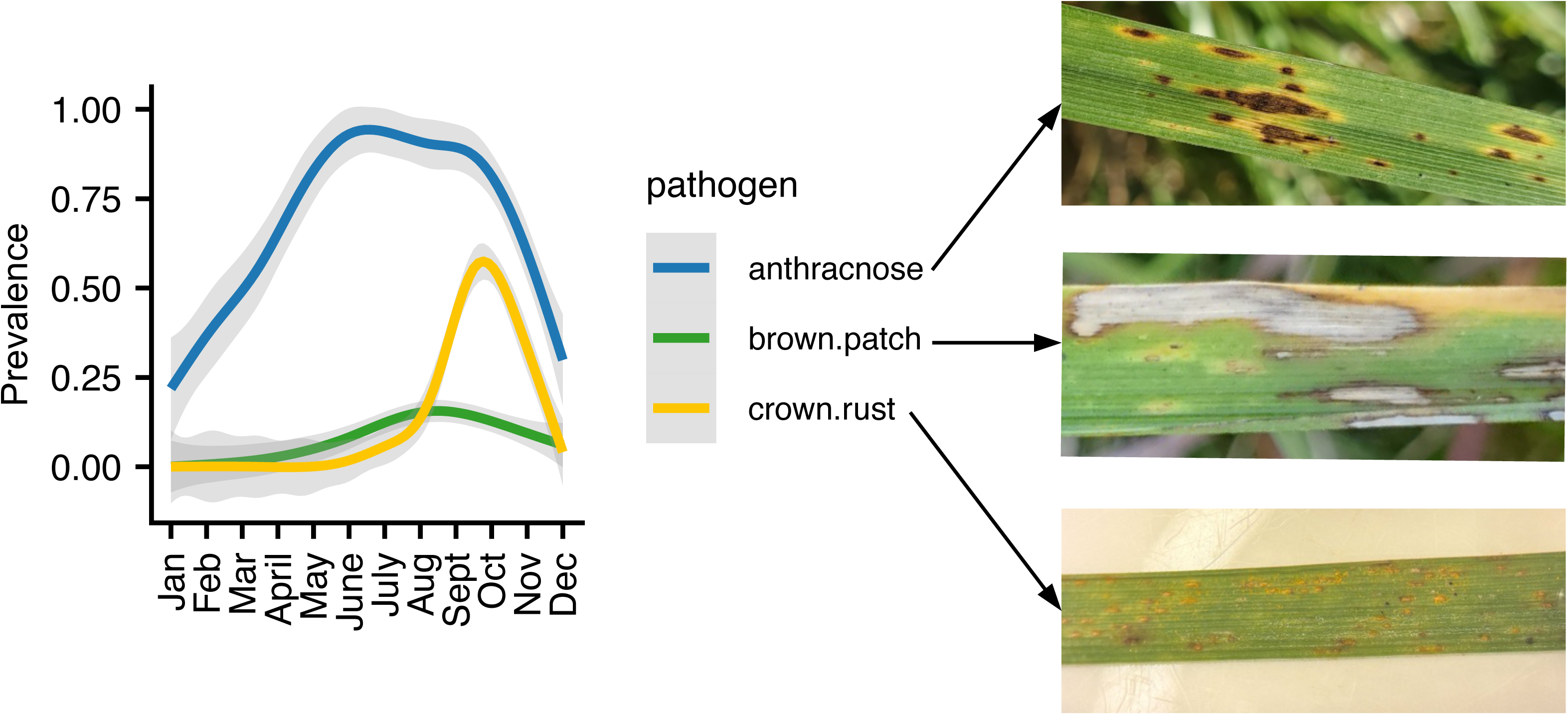
Mean seasonal dynamics for anthracnose, brown patch, and crown rust diseases. For each disease, the trendline represents a cubic spline and 95% confidence interval fit through observations of disease prevalence in the full dataset from 2017 to 2024. Photos at right give an example of each disease on tall fescue. Annual dynamics for each disease can be found in Appendix S1: Figure S1. *Photos: Susan Cleary (top, middle), Dalia Chen (bottom)*.

The three focal diseases exhibit distinct seasonal cycles. Anthracnose prevalence peaks in May, followed by brown patch prevalence peaking in August, and crown rust prevalence peaking in late October (Figure 1, Appendix S1: Figure S1; Grunberg et al., 2023). These temporally staggered outbreaks create opportunities for coinfection and potential pathogen interactions that could influence host performance and disease severity over the growing season.

### Disease Survey Data

All disease surveys were conducted between 2017 and 2024 in plots of previously established old-field vegetation. Plant communities in the plots were typically dominated by tall fescue, and other plant species were not tracked for disease. We combined published disease data (Grunberg et al., 2025) with our own novel data to generate a seven-year dataset of seasonal disease occurrence between 2017 and 2024 (Appendix S1: Table S1).

Each survey in this data set recorded the presence or absence of our three focal diseases at the plant- (i.e., tiller-) or leaf-level. We reduced multiple observations to the plant level, where if disease symptoms were observed on any leaf, we marked the disease as present for the plant. In each survey, experimental plots were tracked through time, yielding plot-level trajectories for all three focal diseases. Not all experimental plots were surveyed for the entire seven-year window. Additionally, some past experiments involved the treatment of experimental field plots with fungicide. For these datasets, only control (i.e. no-fungicide) plots were retained for the population-level Bayesian and random forest analyses. In total across our study, this yielded 29,965 plant-level observations.

In 2018, within some experimental plots, individual plants were tagged and tracked longitudinally through time. These plant-level longitudinal data were used to estimate effects of earlier arriving pathogens on later arriving pathogens via survival analysis (e.g., following Halliday et al. (2017), Halliday et al. (2018), and Halliday et al., (2020); Appendix S1:Table S1).

### Environmental Data

Weather station data was obtained from the North Durham Wastewater Treatment Facility (ID: DURH; North Carolina State Climate Office, 2025). This facility is approximately 10 km from Widener Field and was the closest station which recorded both atmospheric and soil conditions at all time points surveyed for disease. The final data set included 2,814 observations (daily mean value) of 32 climatic variables spanning April 24, 2017, to June 6, 2025. A full list of climatic variables, units, and descriptors is given in Appendix S1: Table S2.

For analyses sensitive to multicollinearity, observations of the 32 climatic variables through time were reduced to four principal components (PCs) using the R function ‘prcomp’. Climatic variables were first centered and scaled to have a mean of zero and variance of one using the ‘decostand’ function in R package vegan (method = “standardization”). The first four PCs together explained 79% of the total variance in the climatic variables over time and explained more variance than expected by chance under a broken stick model (Appendix S1: Figure S2). For the week preceding each fungal disease observation, we calculated the mean score for each principal component. The average dynamics of these environmental PCs over the course of a year can be found in Appendix S1: Figure S3. Finally, we visualized the loadings of each climatic variable on the PCs to infer relationships between the original 32 variables and 4 summary PCs (Appendix S1: Figure S4).

### Random forest analysis to identify abiotic thresholds and predict annual pathogen dynamics

Random forest models were developed using the R packages ‘ranger’ and ‘tidymodels’ (Kuhn & Wickham, 2020; Wright & Ziegler, 2017). We used random forest models to predict disease occurrence from abiotic and biotic conditions, identify the most important predictors, and detect potential thresholds in environmental gradients associated with increased disease risk. This approach also allowed us to accommodate collinearity and nonlinear relationships among predictors. We trained binary classification models to predict the presence or absence of each pathogen at the plant level. Predictor variables included the 7-day average of climatic variables preceding each pathogen observation, the presence or absence of other pathogens on the same plant, the mean prevalence of each pathogen in each plot at the previous time point, month, and year. For example, the crown rust model had the following structure:

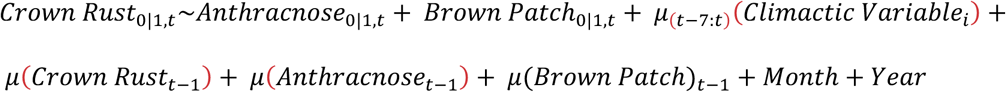

To avoid temporal data leakage (where future information is inadvertently used to train or evaluate a model and thus leads to unrealistically good performance), the data was split into training and test sets based on year and month of observation. The first 80% of the data from May 2017 – July 2022 was used as training sets, whereas the remaining 20% of the data from August 2022 – November 2024 was used as the test set.

Within the training data, we applied time series cross-validation using a sliding window approach. Each validation fold used the prior 12 months of data as training and tested on the subsequent month. This procedure generated a total of 36 sliding periods, though six of these contained validation sets with insufficient data (all spanning winter months, December – February) and were excluded, for a total of 30 final rolling windows. To address class imbalance, the data was up-sampled so that the number of pathogen presence cases equaled the number of absence cases in each temporal split. The total number of observations of each disease across the sampling window are given in Appendix S1: Table S3.

For each random forest model, we grew 3,000 trees. The mtry hyperparameter, which controls the number of variables considered in each decision tree, was optimized via grid search over values from 4 to 32. The value minimizing the out-of-bag (OOB) error was selected for each model. The final ‘mtry’ parameter used for each model was 12 for anthracnose, 8 for crown rust, and 12 for brown patch.

Overall model performance was evaluated by comparing predictions to the held-out test set. We evaluated overall performance using accuracy, recall, precision, F-score, and kappa, which are described further in Appendix S1: Table S4. Variable importance was assessed on the test data and is reported as permutation-based accuracy, indicating accuracy improvement for each predictor when compared to the permuted predictor. The relationship between climatic variables and focal pathogen presence was further explored using accumulated local effects (ALE) plots. We explored the ALE for variables broadly representing conditions important to fungal biology and predictive of disease outcomes in at least one of three focal pathogens. ALE plots facilitate understanding of how individual predictor variables affect the response of a machine learning model, while controlling for cross-correlation and interactions among predictors. In our case, a higher ALE value indicates that the model is more likely to predict pathogen presence for a given parameter value than the model would predict on average.

### Environmental niche overlap between pathogens

Because associations between pathogen species identified in our machine learning models could result from correlated environmental preferences rather than biotic interactions, we additionally quantified overlap between pathogen species’ environmental niches using the R package ‘nicheROVER’ (Lysy et al., 2023). We used the full plot-level dataset for this analysis to evaluate shared environmental preferences as an alternative explanation for the associations between pathogens identified by our models. Climatic variables were summarized as PC scores for this analysis. Using this information, we calculated the pairwise probabilities that a randomly drawn individual from pathogen A would be found within the interior 95% of the niche region of pathogen B using function ‘overlap’ (alpha = .95). *Survival analyses to estimate effects of earlier arriving pathogens on later arriving pathogens*

To better distinguish associations driven by shared environmental preferences from those arising through infection history, we used survival analyses to examine how prior infection influenced subsequent disease risk (e.g., Halliday et al., 2017; Halliday et al., 2018; and Halliday et al., 2020). In 2018, individual plants were surveyed longitudinally (i.e., marked and tracked over time) for disease. While the random forest models utilized only control samples from each field experiment included, we leveraged the treatment groups from the 2018 field experiment to better understand pathogen-pathogen interactions. In this experiment, field plots were either never sprayed with fungicide (control) or sprayed with fungicide for 7-months, 9-months, or continuously to interrupt the onset of brown patch and anthracnose infections. We included control, 7-months, and 9-months treatment groups in the survival analysis. In the control group, out of 432 diseased plants, 14 were infected first by brown patch, 10 were infected first by crown rust, and 408 were infected first by anthracnose. We therefore supplemented these observations with the treatment groups described above, for a total of 1,211 tracked plants, 31 of which were infected first by crown rust, 32 of which were infected first by brown patch, and 1,201 of which were infected first by anthracnose (diseases which arrived at the same timepoint were counted towards the total for both).

Using this data, a survival analysis was conducted for each focal disease to evaluate whether its infection rate was increased or decreased by prior infection by the other two pathogens, in the context of the environment. For each pathogen, we constructed a Cox proportional hazards regression model, with plot ID included as a clustering variable to account for repeated measures of plants within a plot and environmental variables summarized by the first four principal components as covariates. R package ‘survival’ was used to build models (Therneau, 2026).

### Bayesian hierarchical analysis to estimate the interaction between biotic and abiotic variables

To complement the machine learning and survival models, we additionally constructed Bayesian regression models for our data. This Bayesian framework allowed us to account for hierarchical structure and quantify uncertainty in pathogen–pathogen and pathogen–environment interactions using the full plot-level dataset, which could not be easily addressed within the survival or random forest frameworks.

Separate models were built for each focal pathogen (Bernoulli family with logit link) using the R package ‘brms’ (Bürkner, 2021). Fixed effects included coinfecting pathogens, environmental variables summarized by the first four principal components, sampling month and year, and the focal pathogen’s prevalence in the prior census at the same plot. Interaction terms between co-occurring pathogen presences and environmental variables were also included to assess environmentally dependent changes in the strength or direction of pathogen-pathogen interactions. A random effect was included to account for repeated resampling of plots over time. For example, the crown rust model had the following structure:

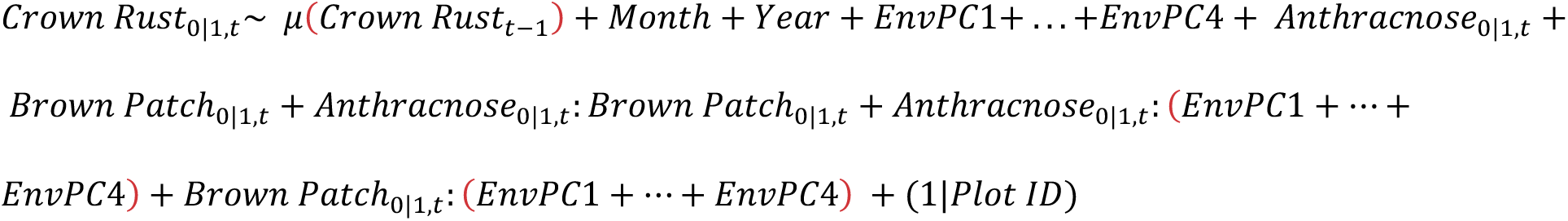

A weakly informative prior, *Normal*(0, 2), was applied to fixed effects to improve convergence and prevent overfitting of terms. Four chains were run per model with 6000 iterations total and 3000 burn-in iterations. Posterior predictive checks and convergence diagnostics (*R*^"^ < 1.01, ESS > 1000) confirmed adequate fit across all models (Appendix S1: Figures S4 – S7).

All data and code used to generate the results contained in this paper are available on GitHub (https://github.com/cb-scott/WidenerLongTermAnalysis) and archived on Zenodo at 10.5281/zenodo.22285642.

## Results

### Predictive accuracy of temporal random forest models for crown rust, brown patch, and anthracnose

Crown rust, anthracnose, and brown patch dynamics were each best predicted by disparate environmental and biotic factors (Appendix S1: Figures S8 – S10). Model performance metrics were comparable across brown patch (Accuracy = 0.937, Kappa = 0.364) and crown rust models (Accuracy = 0.853, Kappa = 0.437), indicating high predictive power. Conversely, the anthracnose model had lower predictive power (Accuracy = 0.555, Kappa = 0.113) and was only nominally better than selecting classes based on random chance (Appendix S1: Table S4). For each disease, we also constructed models with the 14-day and 30-day averages of environmental predictors leading up to the sampling date, rather than the 7-day average presented here. Models with these longer-term averages generally decreased model performance (Appendix S1: Tables S5 – S6). Visualizations of the top 15 most important variables (after this threshold there is a generally steep drop off in variable importance and most variables are colinear) are given in Appendix S1: Figures S11 – S13.

### Species-specific environmental thresholds and biotic interactions identified from random forest analysis

Crown rust dynamics were primarily driven by moisture-, light-, and wind-related variables, as well as anthracnose and brown patch coinfection and their lagged prevalences (Appendix S1: Figure S8). However, the best predictor for crown rust presence on a host plant was its own lagged prevalence, representing disease inoculum. Beyond this, crown rust was most likely to be observed when total precipitation was low, maximum photosynthetically active radiation ranged from 1000-2000 μmol/m^2^/s, maximum daily relative humidity exceeded 98%, and average wind speeds were less than 4 or greater than 6 kmph (Figure 2a-d; Appendix S1: Figure S11). Crown rust presence at the plant level was also positively associated with anthracnose coinfection but negatively associated with brown patch coinfection (Figure 2e).

**Figure 2.**
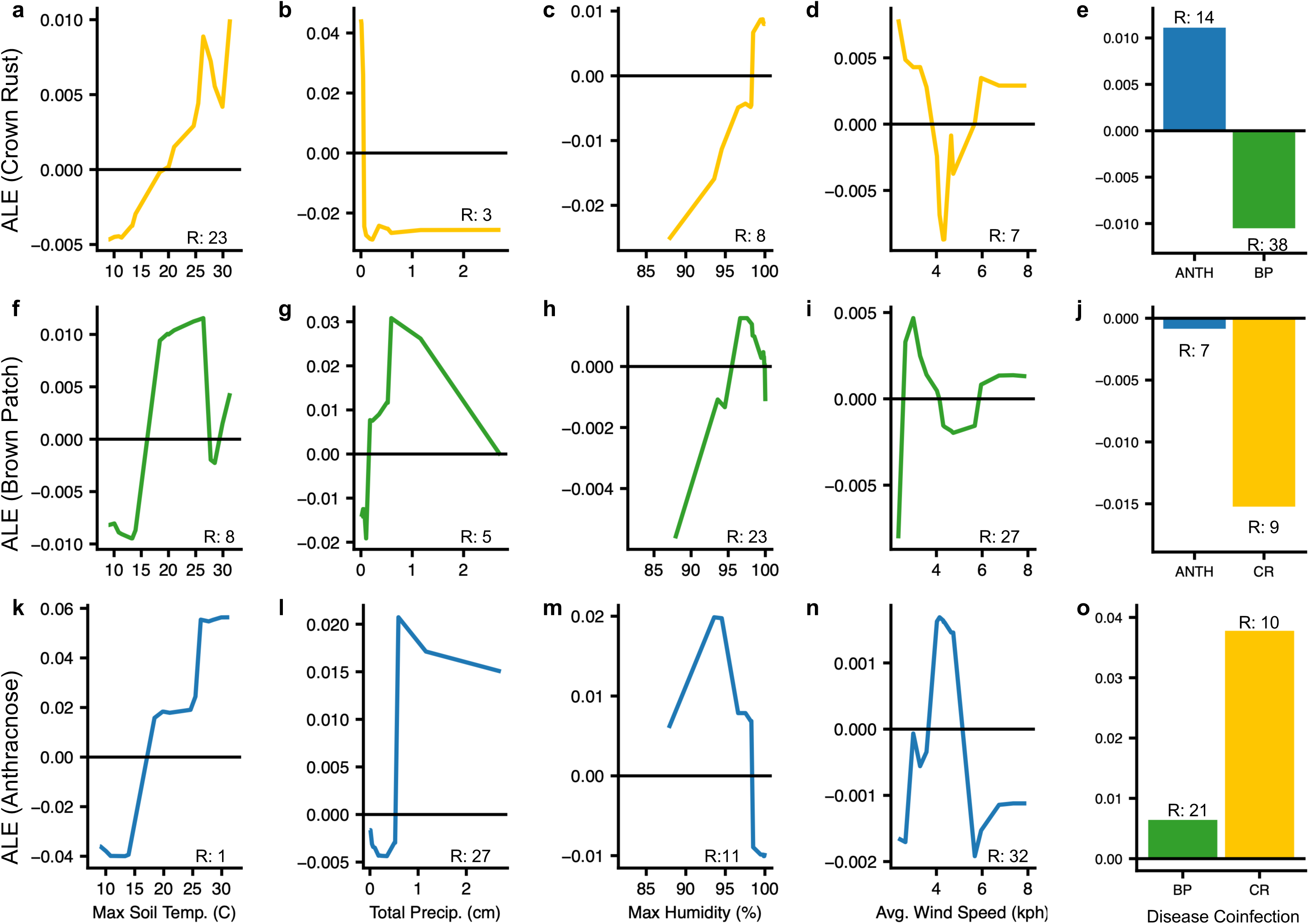
Random forest models identified key environmental and biotic predictors associated with disease dynamics, including temperature, precipitation, humidity, wind speed, and disease coinfection. Each plot shows the accumulated local effects (ALE) for a single predictor included in the focal disease’s model. ALE magnitude coarsely scales with variable importance, so larger ALE values generally reflect greater predictive value when all other variables are held at their mean. Positive ALE values indicate increased occurrence of the focal disease; negative values indicate decreased occurrence. "R:" gives each variable’s importance rank out of the 39 variables included in the model. Steep changes in ALE, especially where the sign changes, mark putative environmental thresholds. (a-e) Crown rust. (f-j) Brown patch. (k-o) Anthracnose. The same variables are shown for each disease to allow direct comparison. All variables and their ALEs are given in Appendix S1: Figures S11-S13.

Brown patch dynamics were primarily predicted by rainfall- and temperature-related variables and lagged anthracnose and crown rust prevalence (Appendix S1: Figure S9). Like crown rust, the best predictor of brown patch presence at the plant level was its own lagged prevalence in the host population, representing disease inoculum. With respect to the environmental variables, brown patch was most likely to be observed when daily precipitation ranged from 0.5-2.5 cm and when maximum soil temperature ranged from 15-27.5°C (Figure 2f-i; Appendix S1: Figure S12). Anthracnose or crown rust coinfection decreased the probability of observing brown patch, and this effect was strongest for crown rust (Figure 2j).

While overall predictive power for anthracnose was lower, its dynamics were primarily driven by temperature-, moisture- and humidity-related variables (Appendix S1: Figure S10). Anthracnose was most likely to be observed at the plant-level when daily average soil temperatures exceeded 17°C, daily average soil moisture was lower than 0.44 m^3^/m^3^, and daily maximum relative humidity ranged from 87.5-98% (Figure 2k-n; Appendix S1: Figure S13). While neither crown rust nor brown patch coinfection, or their previous prevalences in the host population, were among the most important variables in predicting anthracnose disease presence, crown rust coinfection increased the probability of observing anthracnose more than brown patch (Figure 2o).

### Environmental niche overlap among pathogens

Environmental niche analyses using the full dataset revealed substantial climatic niche overlap among anthracnose, brown patch, and crown rust(Appendix S1: Figure S14), making it possible that the apparent facilitative effects identified by our machine learning models result from shared environmental responses rather than biotic interactions. Moreover, these potential interactions may be masked in the full combined dataset from 2017–2024, since it contains repeated measurements at the plot level but lacks longitudinal tracking of individual plants in most years.

### Survival analyses of plant-level disease risk

To further investigate the role of historical contingencies in disease risk, we utilized a subset of the dataset from 2018 which contained longitudinal, individual plant-level data to test how prior pathogen infections and environmental variables influence the host individual’s risk of acquiring each disease (“disease risk” for a focal disease) using Cox proportional hazards models.

The full crown rust model provided a significantly better fit than the null model (LRT: χ² = 604.5, df = 6, *p* < 0.001; Harrell’s C = 0.738 ± 0.021), with climatic variables and prior infection by anthracnose emerging as significant predictors of disease risk (Figure 3a). Prior infection by anthracnose increased the risk of subsequent crown rust infection approximately 2-fold (Hazard ratio: 2.18, *p <* 0.01). Prior infection by brown patch had no significant effect on crown rust infection (Hazard ratio: 1.28, *p* = 0.081). Three of the four environmental PCs significantly affected the risk of crown rust infection, albeit with smaller effect than prior infection by anthracnose. The inclusion of prior infection status significantly improved model fit beyond environmental PCs alone (LRT: χ² = 16.4, df = 2, *p <* 0.01).

**Figure 3.**
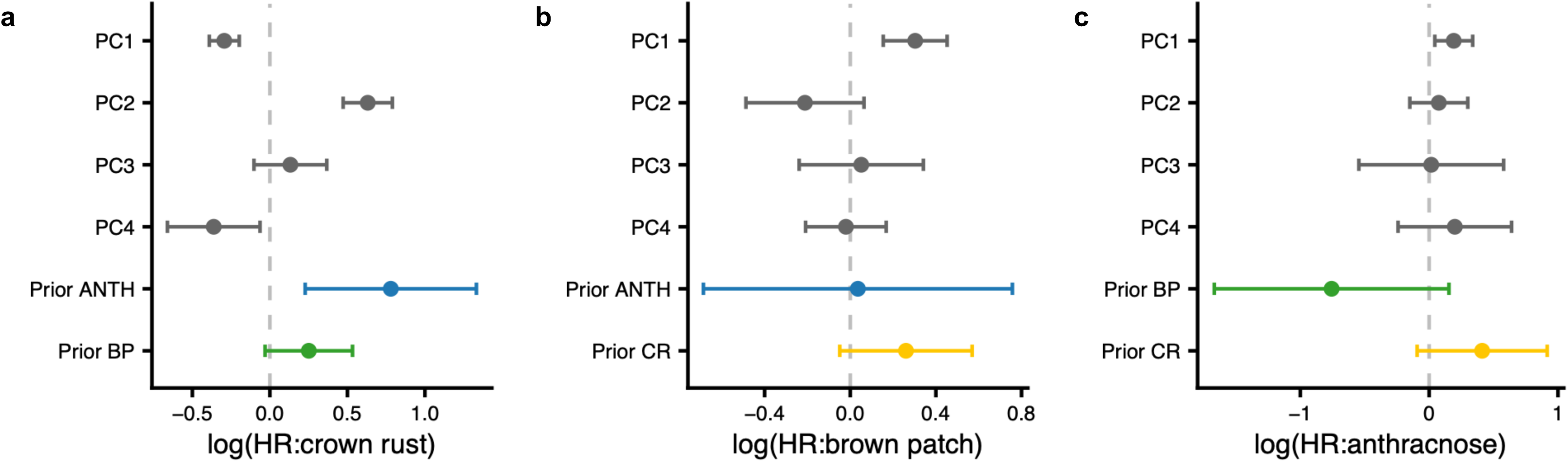
Survival models revealed that the strongest predictor of crown rust disease was prior infection by anthracnose, and that all three diseases were sensitive to environmental conditions. Models were built from the 2018 longitudinal data, in which field plots were perturbed with fungicide to manipulate disease arrival timing. (a) Crown rust infection risk increased with prior anthracnose infection, decreased in PC1 and PC4, and increased in PC2. Brown patch (b) and anthracnose (c) risk, by contrast, increased only with increases in PC1, which primarily represented warmer temperatures.

For brown patch, the full model also provided a significantly better fit than the null survival model (LRT: χ² = 143.8, df = 6, p < 0.001; Harrell’s C = 0.644 ± 0.027). Neither prior infection by anthracnose (Hazard ratio: 1.03, *p* = 0.923) nor crown rust (Hazard ratio: 1.29*, p* = 0.099) significantly affected the risk of brown patch infection (Figure 3b). Similarly, the inclusion of these predictors did not improve model fit beyond environmental PCs alone (LRT: χ² = 4.21, df = 2, *p =* 0.122). Instead, environmental PC1 had the largest, and the only significant, effect on brown patch risk (Hazard ratio: 1.35, *p* < 0.01), likely representing the effect of temperature (Appendix S1: Figure S4).

Finally, for anthracnose, the full model did not provide a significantly better fit than the null survival model, though concordance was similar to rust and brown patch models (LRT: χ² = 11.88, df = 6, *p =* 0.065; Harrell’s C = 0.642 ± 0.040). Neither prior infection by brown patch (Hazard ratio: 0.469, *p* = 0.103) nor crown rust (Hazard ratio: 1.509, *p* = 0.110) significantly affected the risk of anthracnose infection (Figure 3c). The inclusion of these predictors did not improve model fit beyond environmental PCs alone (LRT: χ² = 3.09, df = 2, *p =* 0.214). However, environmental PC1 had a significant and positive effect on anthracnose risk (Hazard ratio: 1.21, *p* < 0.05), again likely representing the effects of higher temperature (Appendix S1: Figure S4).

### Bayesian hierarchical analysis of context-dependent associations between pathogens

Finally, we investigated potential interactions between the four environmental PCs and pathogen co-occurrence in driving focal disease presence. We built a Bayesian hierarchical model to predict the probability of observing the focal pathogen based on environmental PCs, time of year, pathogen coinfection, and the two-way interactions of PCs and coinfection. By using these models, we were able to explicitly account for the repeated sampling of plots through time and retained sufficient power to look at interactions between environmental PCs and pathogen coinfection, lending us insights into how pathogen facilitation may strengthen or weaken under specific environmental contexts. For each disease, we found credibly non-zero interactions between environmental PCs and pathogen coinfection. Here, we focus on these interaction terms, visualizing them along the temperature (PC1) gradient, as they give insights not revealed by our other models. Full Bayesian model results can be found in Appendix S1: Tables S7 – S9 and all conditional effects are visualized in Appendix S1: Figures S15-S17.

For crown rust, coinfection by either anthracnose or brown patch interacted with the effects of the abiotic predictors for most environmental PCs (Appendix S1: Table S7). The variable which was most predictive of crown rust presence was anthracnose coinfection (*β* = 1.15, *CI*_95%_ = [0.976, 1.33]). Increases in PC1, characterized by higher temperatures, increased the probability of crown rust infection (*β* = 0.132, *CI*_95%_ = [0.0717, 0.193]), and this association was amplified in plants coinfected by brown patch (*β* = 0.146, *CI*_95%_ = [0.0812, 0.181]) and reduced for those coinfected by anthracnose (*β* = −0.051, *CI*_95%_ = [−0.0884, −0.0131]; Figure 4a,b). Increases along PC2, characterized by higher humidity, similarly increased crown rust infection probability (*β* = 0.123, *CI*_95%_ = [0.0213,0.225]), but this effect was generally reduced for plants coinfected by anthracnose (*β* = −0.0704, *CI*_95%_ = [−0.168, 0.0329]) or brown patch (*β* = −0.205, *CI*_95%_ = [−0.315, −0.0936]). Increases along PC3, which was not well characterized by any individual environmental variable, generally did not affect the probability of crown rust infection (*β* = −0.240, *CI*_95%_ = [−0.149, 0.0987]). Finally, increases in PC4, characterized by decreasing soil moisture, decreased the probability of crown rust infection (*β* = −0.302, *CI*_95%_ = [−0.393, −0.214]). For plants coinfected by either brown patch (*β* = 0.0757, *CI*_95%_ = [−0.0257, 0.181]) or anthracnose (*β* = 0.114, *CI*_95%_ = [0.0239, 0.200]), this negative relationship was generally more pronounced.

**Figure 4.**
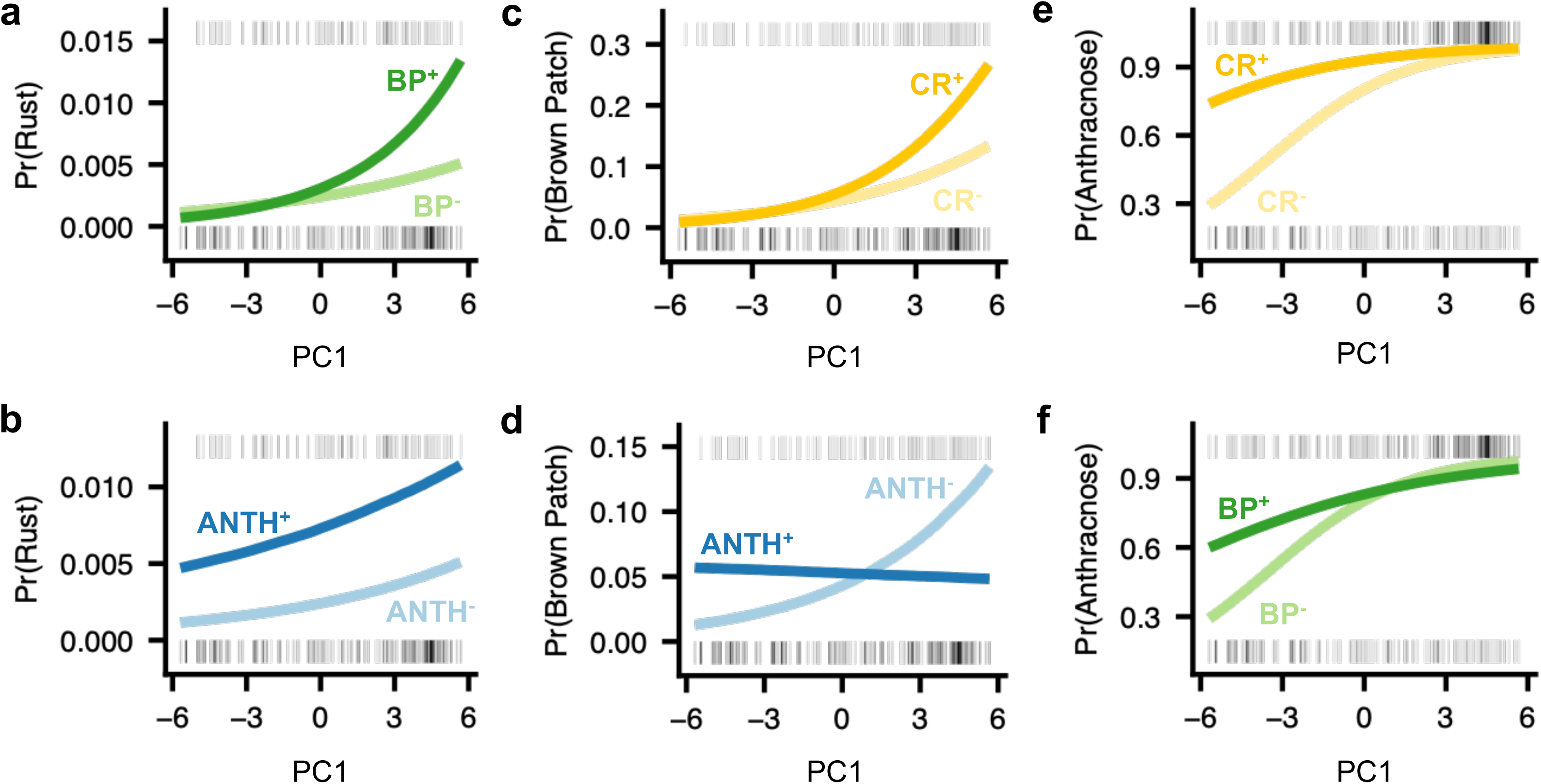
Coinfection can modulate the effects of environmental gradients on disease risk. Panels show conditional effects of PC1 × co-infection interactions from the Bayesian hierarchical models. Darker lines represent disease presence; ANTH^+^/ANTH^−^ denotes presence/absence of anthracnose, CR^+^/CR^−^ denotes presence/absence of crown rust, and BP^+^/BP^−^ denotes presence/absence of brown patch. Rugs along the top of each inset give the density of focal disease presences along the environmental PC; rugs along the bottom give the density of focal disease absences. Response variables are crown rust (a-b), brown patch (c-d), and anthracnose (e-f). All interactions shown had non-zero posterior credible intervals. Conditional effects of all pathogen × PC interactions are shown in Appendix S1: Figures S15-S17.

For brown patch, coinfection by either anthracnose or crown rust interacted with environmental PC1 and PC2 (Appendix S1: Table S8). Higher PC1 values, representing higher temperatures, increased the probability of brown patch infection (*β* = 0.219, *CI*_95%_ = [0.149, 0.287]), and this effect was intensified by crown rust coinfection (*β* = 0.107, *CI*_95%_ = [0.0594, 0.156]; Figure 4c). Conversely, coinfection by anthracnose decreased the probability of brown patch infection at high PC1 values (*β* = −0.223, *CI*_95%_ = [−0.280, −0.189]; Figure 4d). Higher PC2 values, representing increased humidity or lower winds, also increased the probability of brown patch infection (*β* = 0.327, *CI*_95%_ = [0.225, 0.432]). Coinfection by either anthracnose (*β* = −0.280, *CI*_95%_ = [−0.382, −0.184]) or crown rust (*β* = −0.190, *CI*_95%_ = [−0.292, −0.109]) lessened this effect, decreasing the probability of brown patch infection at higher PC2 values. While higher PC3 (*β* = 0.135, *CI*_95%_ = [0.00399, 0.261]) and lower PC4 (*β* = −0.0658, *CI*_95%_ = [−0.160, 0.028]) values likely increased the probability of brown patch infection, they did not interact strongly with anthracnose and crown rust coinfection.

Finally, for anthracnose, coinfection by crown rust or brown patch frequently influenced the association between abiotic predictors and anthracnose infection (Appendix S1: Table S9). Higher environmental PC1 values, indicating higher temperatures, were positively associated with anthracnose infection (*β* = 0.403, *CI*_95%_ = [0.361, 0.445]). Plants with crown rust (*β* = −0.132, *CI*_95%_ = [−0.172, −0.0914]) or brown patch coinfections (*β* = −0.194, *CI*_95%_ = [−0.241, −0.146]) were less tightly associated with changes along PC1 (Figure 4e,f). PC2 was positively associated with anthracnose infection (*β* = 0.120, *CI*_95%_ = [0.0794, 0.162]), where high values generally represent high humidity. Brown patch coinfection reduced the effects of PC2 on anthracnose infection (*β* = −0.227, *CI*_95%_ = [−0.327, −0.127]). Increased PC3 values, which likely represents a combination of wind, moisture, and sunlight variables, also increased the probability of anthracnose infection (*β* = 0.334, *CI*_95%_ = [0.279, 0.388]). For plants coinfected by brown patch, this effect was lessened (*β* = −0.254, *CI*_95%_ = [−0.377, −0.131]), while coinfection by crown rust intensified this effect (*β* = 0.151, *CI*_95%_ = [0.0365, 0.271]). Finally, increases in PC4, likely representing decreased soil moisture, decreased the probability of anthracnose infection (*β* = −0.216, *CI*_95%_ = [−0.270, −0.163]). Brown patch (*β* = 0.143, *CI*_95%_ = [0.0519, 0.236]) or crown rust (*β* = 0.0987, *CI*_95%_ = [0.014, 0.180]) coinfection both lessened the negative effects of PC4 on anthracnose infection probability.

## Discussion

These results have several implications, first ecological, and then methodological. Our analysis of multiple pathogens across seven years revealed that, while seasonal epidemics are fundamentally driven by changes in abiotic conditions, disease risk was often also predicted by pathogen coinfections. For crown rust, the influence of pathogen coinfection, particularly facilitation by anthracnose, rivaled that of abiotic conditions. Furthermore, our results suggest that the strength of coinfections’ influence varied with environmental context and that coinfections can modulate the effects of abiotic drivers on seasonal epidemics.

Methodologically, temporally explicit random forest models generated accurate predictions of field epidemics for two of three focal diseases and identified critical environmental thresholds for all three, illustrating the predictive power of these models. We then combined these machine learning models with Bayesian hierarchical models and survival analysis models to add mechanistic understanding. In our study of seasonal epidemics in a multi-pathogen system, employing multiple complementary models provided an integrated framework for ecological understanding and forecasting. This framework may be valuable for studies in other systems in resolving the roles of diverse ecological processes that vary over time.

Past field studies in this system have documented species interaction networks including both antagonistic and facilitative interactions between pathogens, and that these interactions varied over a single growing season (Grunberg, Joyner, et al., 2023; Halliday et al., 2017). Additionally, past work in the system has established the importance of seasonal environmental conditions to interannual variation in pathogen community assembly (Grunberg et al., 2026). Here, analyzing seven years of data using Bayesian hierarchical models allowed us to explicitly examine how pathogen species interactions are influenced by specific seasonal environmental conditions. These models indicated that pathogen species interactions commonly modulated effects of environmental variation on disease. This was particularly true along PC1, representing temperature, with some pathogen interactions estimated to be stronger at higher temperatures and others stronger at lower temperatures. Notably, coinfection by anthracnose increased the probability of brown patch infection at low PC1 values, potentially consistent with the environmental conditions of lab experiments in which inoculation of plants with *Colletotrichum* (i.e. anthracnose) facilitated *Rhizoctonia* (i.e. brown patch; Geyer et al., 2026; Green et al., 2024; O’Keeffe et al., 2021). Such environmentally dependent interactions may be particularly consequential if they broaden the realized environmental niche of a pathogen, enabling infection outside the windows predicted from abiotic conditions alone.

Facilitation was most pronounced for crown rust, where both prior infection by anthracnose, and brown patch to a lesser extent, increased infection probability. Given that *P. coronata* obligately feeds on living tissue (i.e., is an obligate biotroph) while *R. solani* and *C. cereale* both have necrotrophic stages that reduce living host tissue, we initially expected the opposite pattern. This facilitation may instead reflect antagonism between immune-signaling pathways, which is a known source of facilitation among symbiotic organisms in many host-associated systems (Halliday et al., 2024) and can also impact within-host interactions in this system (Halliday et al., 2018; but see Geyer et al., 2026). Specifically, the salicylic- and jasmonic-acid mediated defense pathways are mutually antagonistic in plants, and necrotrophic infection can suppress SA-dependent resistance that would otherwise limit biotrophs like *P. coronata* (Pieterse et al., 2012).

Alternatively, facilitation between any pair of pathogens may reflect underlying variation in host susceptibility rather than a direct pathogen-pathogen interaction where hosts readily infected by early-season pathogens may simply be less resistant overall, and thus more susceptible to later-season infections as well (Man et al., 2023). These context-dependent interactions could additionally reflect shared seasonal phenology (Kumata & Sasaki, 2026). This challenge of distinguishing direct interaction effects from temporal co-occurrence motivated our inclusion of month as a covariate in the Bayesian and random forest models, allowing us to assess the predictive power added by including environmental PCs as well as time of season per se (i.e. month). In some models, month was a stronger predictor than environmental PCs, reaffirming that seasonal timing, such as the arrival of crown rust from alternate hosts or shifts in plant community composition, plays a substantial role in disease emergence and persistence. Nonetheless, in all models, environmental PCs still had non-zero effects after the inclusion of time of season.

Thus, accurate prediction of field disease dynamics may require explicitly accounting for both coinfection dynamics and host infection history. When multiple pathogens circulate simultaneously, single-pathogen frameworks can be qualitatively misleading because they can fail to capture effects of pathogen–pathogen interactions (Dutt et al., 2022; Garin et al., 2018; Strauss et al., 2021; Susi et al., 2015). Our results suggest that some pathogens can increase host susceptibility to infection by other, later-arriving pathogens and thereby alter epidemic trajectories, consistent with other evidence that historical contingency can shape disease outcomes (Halliday et al., 2020). In the future, characterization of coinfection and infection history at the individual scale may better clarify these processes.

Beyond pathogens, these results may inform dynamics across symbiotic systems. Historical contingency is likely common in host-associated microbial communities more broadly (Debray et al., 2022), and our results further suggest the strength of these effects is often environmentally dependent rather than fixed. Thus, not only does it matter whether symbionts interact, but also when and under which conditions. In the future, the collection of longitudinal, paired environmental and symbiont data across host organisms will aid in uncovering general rules for context-dependent interactions.

Methodologically, building on prior single-pathogen work (Alves et al., 2025; Shah et al., 2023), we found random forest models to be a robust framework for capturing the nonlinearity and context-dependence introduced by coinfection. They generated robust predictions (accuracy >80%, Kappa > .3) for two of three focal pathogens and recovered biologically relevant environmental thresholds consistent with known drivers of fungal phenology in general (e.g., temperature, moisture; Delmas et al., 2024) and with the specific life cycles of our focal diseases (Butler & Kerns, 2019a, 2019b, 2019c). Further, the comparatively weak model accuracy for anthracnose may represent the pathogen’s biology rather than an inherent failure of the method. As one of the earliest pathogens to arrive in the field each season, anthracnose has few or no established coinfections to draw on as predictors, and its presence may depend more on host density and growth stage than on environmental conditions or pathogen co-occurrence.

However, random forests excel at prediction while often underutilizing experimental design details that could offer deeper mechanistic insight. We therefore complemented this approach with survival models, which leveraged longitudinal plant-level data to further distinguish facilitation from shared environmental niche space. Combining results across all three modeling approaches, random forest, survival, and Bayesian, we inferred key environmental drivers, pathogen-pathogen interactions, and environment-by-pathogen interactions which may drive disease dynamics in our system (Figure 5).

**Figure 5.**
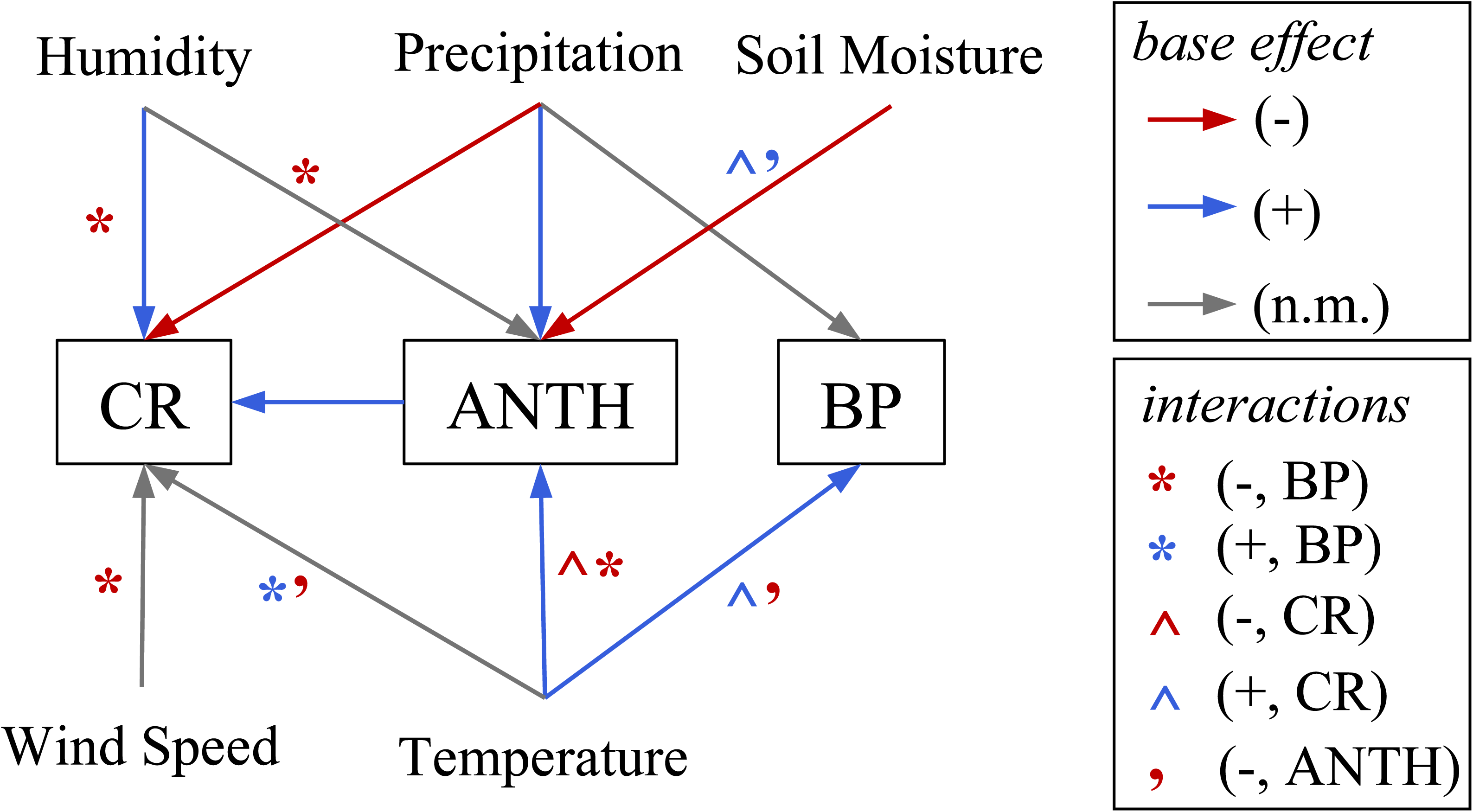
Inferred environmental drivers and biotic interactions for crown rust (CR), anthracnose (ANTH), and brown patch (BP), based on consensus across the three models. For each variable, we report the consensus direction of effect (positive, negative, or non-monotonic) on each pathogen. Asterisks/annotations indicate inferred directions of coinfection-environment interactions.

In the future, ensemble approaches, which formally combine predictions from multiple models, may offer a strategy for modeling multi-pathogen systems (Chauvin et al., 2025; Mao et al., 2025) by pairing near-term machine learning predictions with scenario-based mechanistic forecasts (Ye et al., 2025). This approach may be most powerful when pairing finer-scale environmental measurements (e.g., at the plot level) with long-term, plant-level longitudinal data, enabling integrated forecasts which balance predictive accuracy with mechanistic understanding of how coinfection dynamics shift across environmental gradients. Developing such forecasting capacity will be critical for proactive disease management as climate change alters both pathogen phenology and the environmental context in which pathogen interactions occur.

## Acknowledgements

This project would not have been possible without the many technicians who contributed to fieldwork from 2017-2024. This study was supported by NSF-USDA grant 2016-67013-25762 and NSF grant DEB-2308472 to CEM.

