## Appendix S1 for "Environment and host infection history jointly predict disease risk in a multi-pathogen system"

### Appendix S1: Supplemental Information

Environment and host infection history jointly predict disease risk in a multi-pathogen system

*Ecology*

Carly B. Scott, Susan Cleary, Rita L. Grunberg, Fletcher W. Halliday, Brooklynn N. Joyner, Kayleigh R. O'Keeffe, Isabelle Stiver, Charles E. Mitchell

**References 29**

**Supplemental Figures**

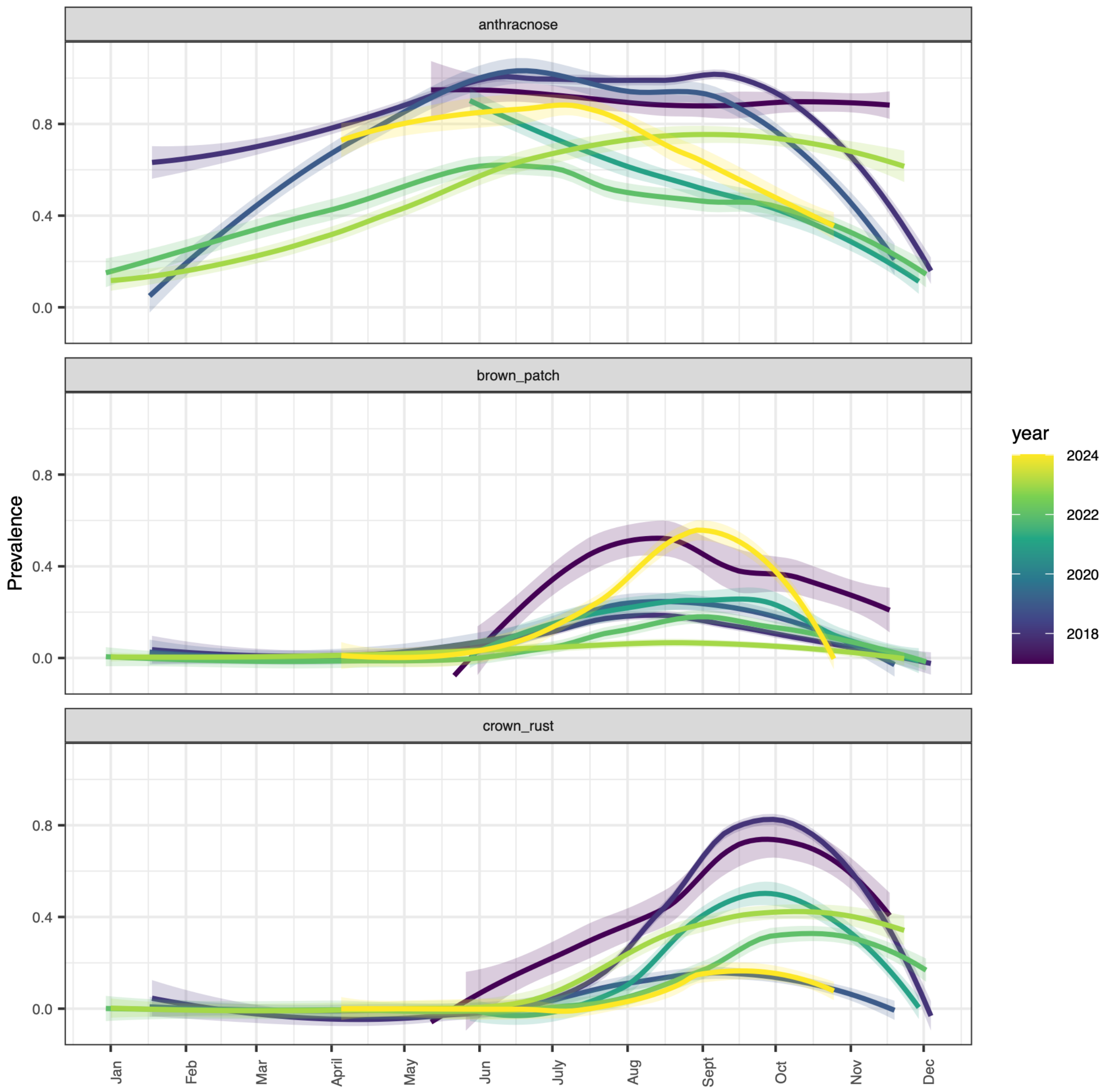

Figure S1. Annual epidemic curves in prevalence of each focal disease (proportion of diseased hosts per plot).

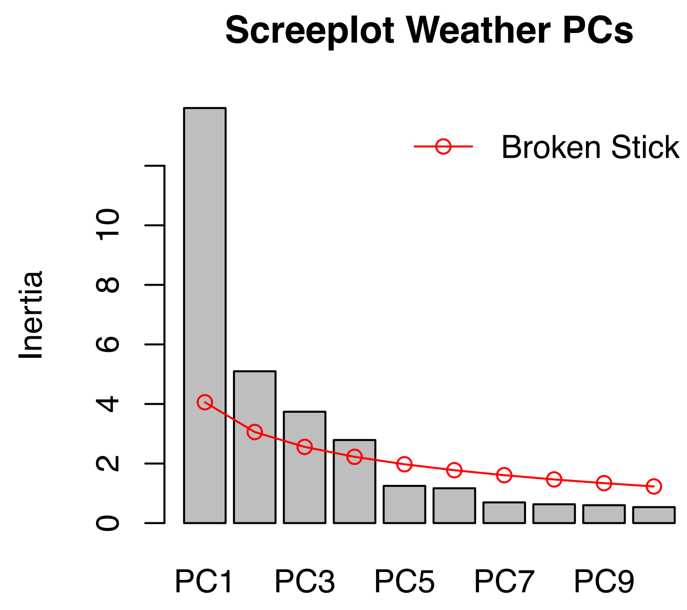

Figure S2. Scree plot of environmental PCs. Red line gives the broken stick null model (amount of variation expected to be explained by chance for each PC). The first four PCs explain more variation than expected by chance and were retained for subsequent analyses.

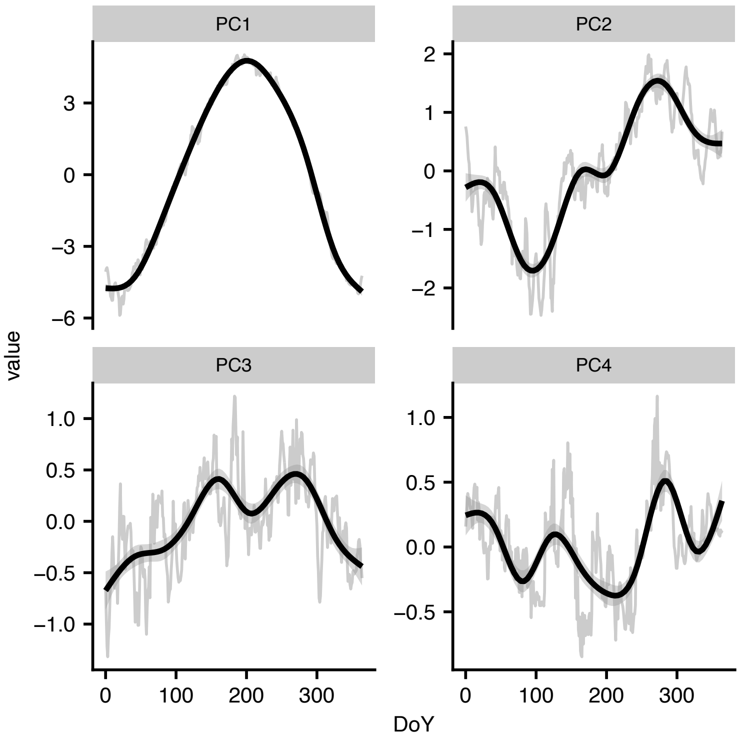

Figure S3. Average dynamics of each environmental PC over the course of a year. DoY represents “Day of the Year” on the x-axis, with day 0 = January 1. Trendline is a general additive model fit to the data with a cubic regression spline and formula y ~ s(x, bs = “cs”).

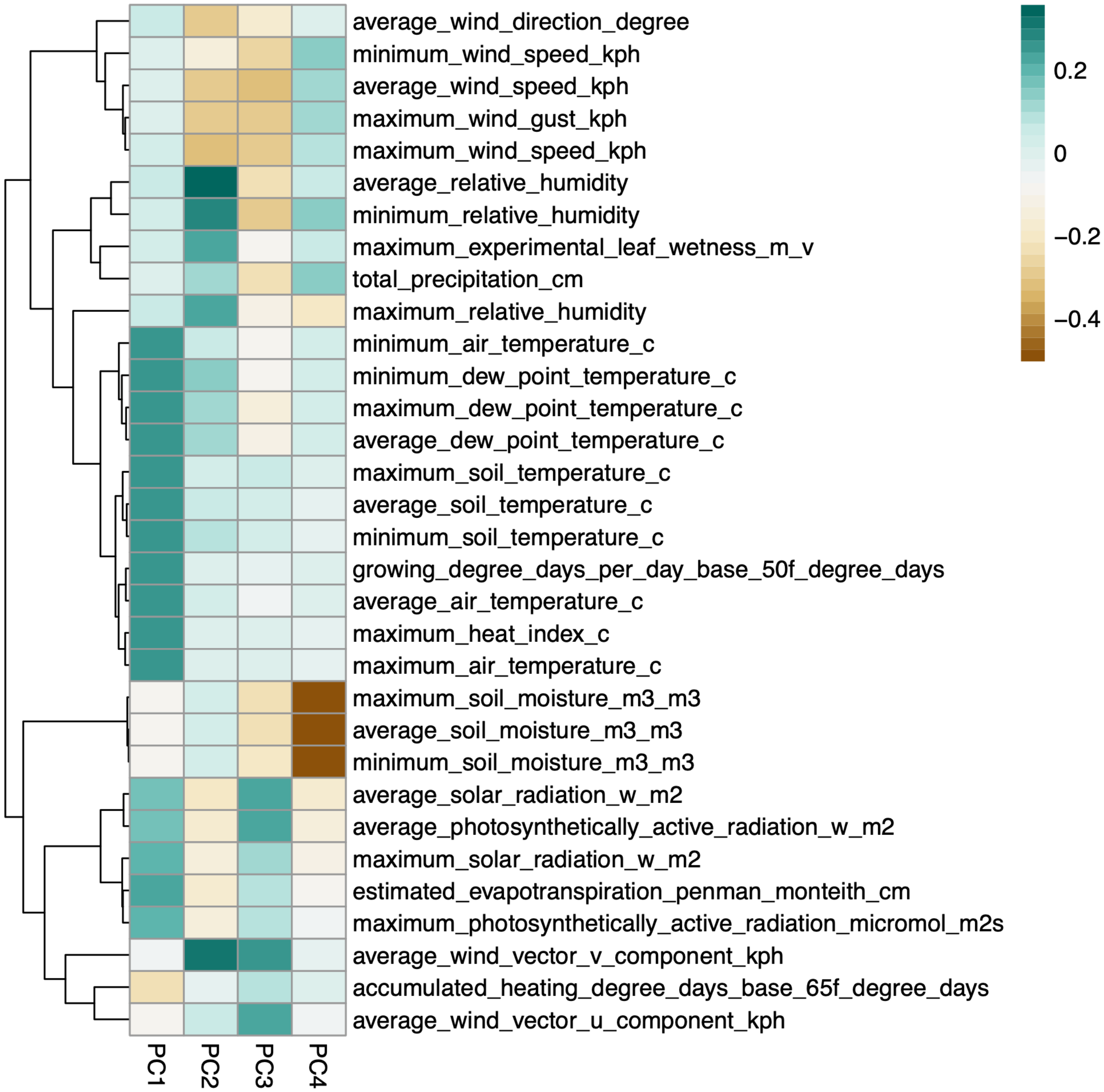

Figure S4. Heatmap of climatic variable loadings on the first four principal components, representing each variable's contribution to the components. Rows are clustered to highlight variables with similar loading patterns across components. Color intensity and hue indicate the magnitude and direction of each loading.

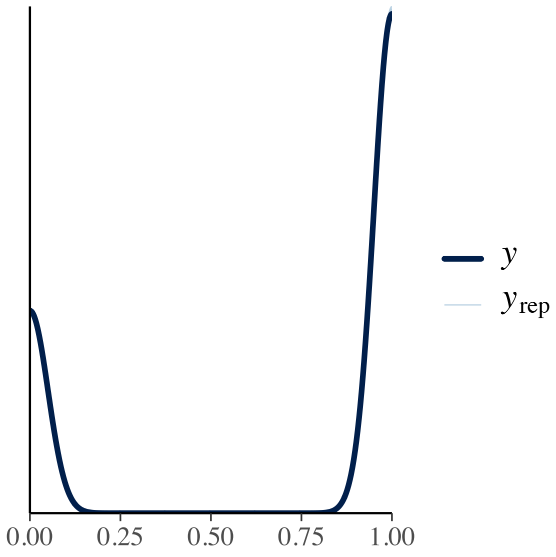

Figure S5. Posterior predictive check for anthracnose ‘brms’ model using a density overlay from 50 posterior draws. The thick line shows the density of the observed data, while the thin, semi-transparent lines represent densities of replicated datasets simulated from the posterior predictive distribution. Good agreement between observed and simulated densities indicates that the model adequately captures the overall distribution of the data; here, thin blue lines are difficult to see as they almost completely overlap with expected values.

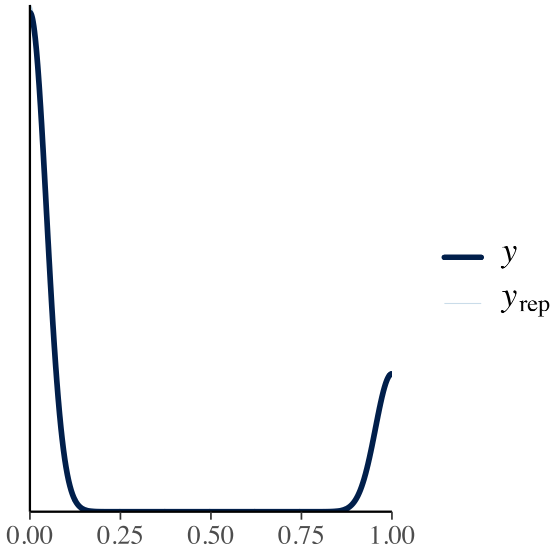

Figure S6. Posterior predictive check for crown rust ‘brms’ model using a density overlay from 50 posterior draws. The thick line shows the density of the observed data, while the thin, semi-transparent lines represent densities of replicated datasets simulated from the posterior predictive distribution. Good agreement between observed and simulated densities indicates that the model adequately captures the overall distribution of the data; here, thin blue lines are difficult to see as they almost completely overlap with expected values.

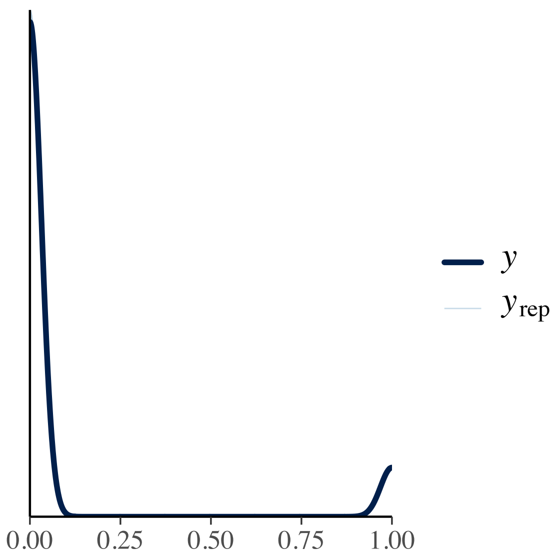

Figure S7. Posterior predictive check for brown patch ‘brms’ model using a density overlay from 50 posterior draws. The thick line shows the density of the observed data, while the thin, semi-transparent lines represent densities of replicated datasets simulated from the posterior predictive distribution. Good agreement between observed and simulated densities indicates that the model adequately captures the overall distribution of the data; here, thin blue lines are difficult to see as they almost completely overlap with expected values.

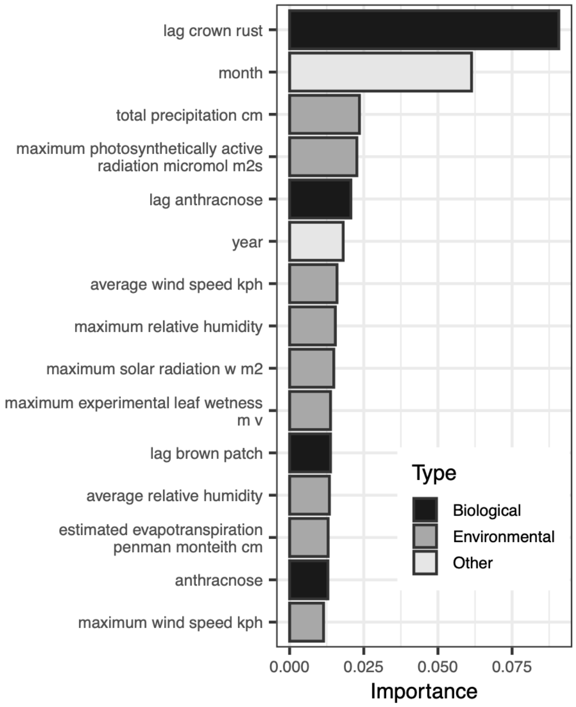

Figure S8. Top fifteen most important variables in predicting crown rust dynamics from random forest model.

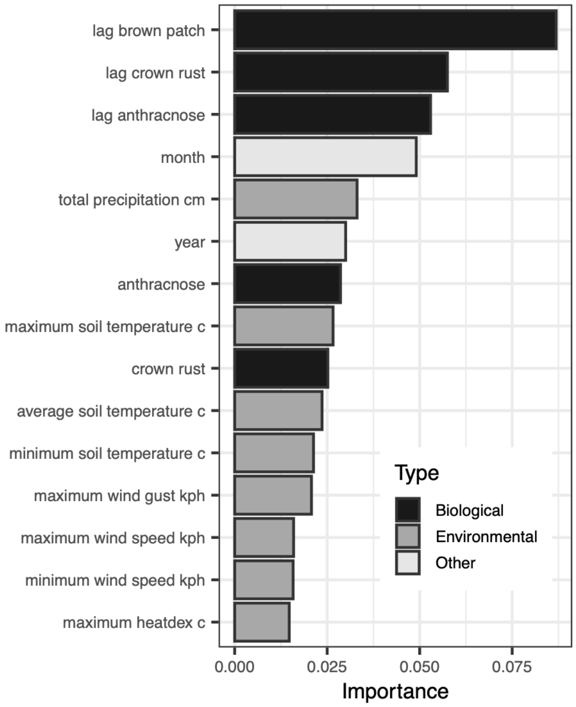

Figure S9. Top fifteen most important variables in predicting brown patch disease state from random forest model.

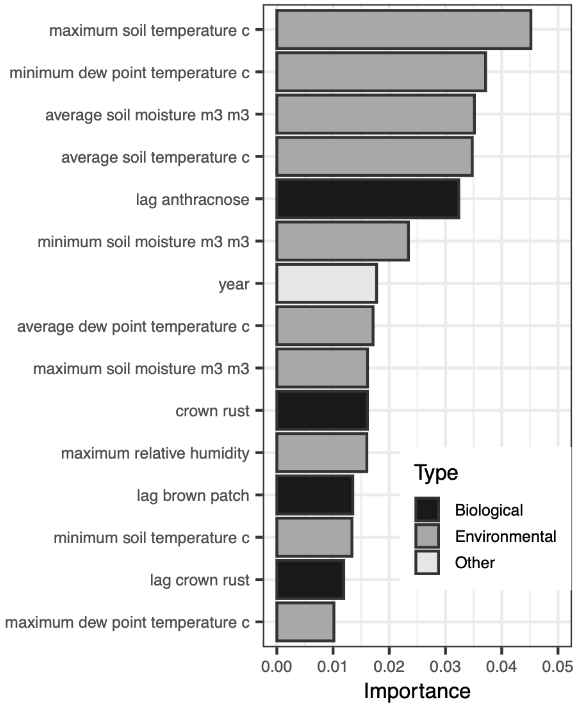

Figure S10. Top fifteen most important variables to predict anthracnose disease state from random forest model.

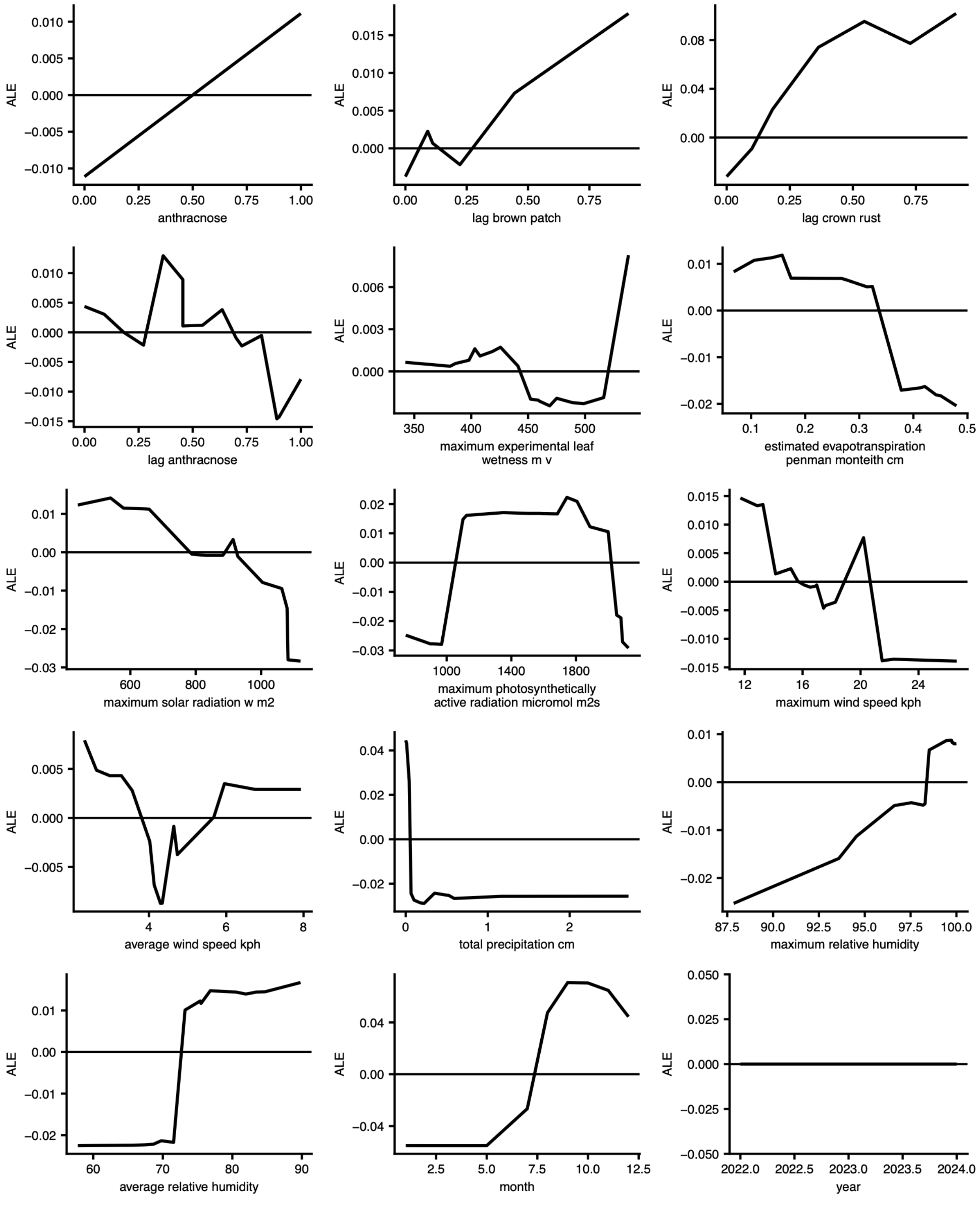

Figure S11. ALE plots for top 15 most important variables in crown rust random forest model.

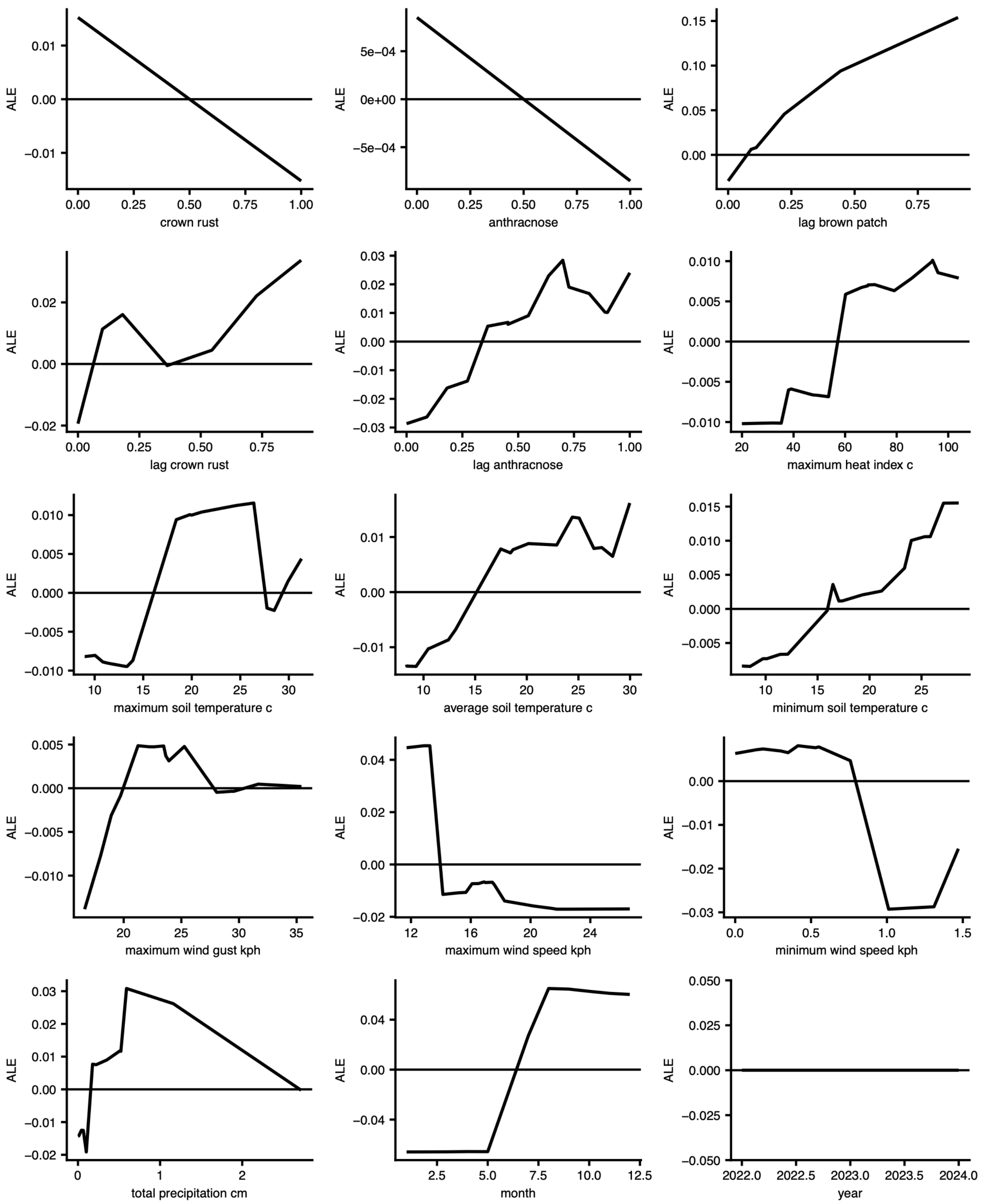

Figure S12. ALE plots for top 15 most important variables in brown patch random forest model.

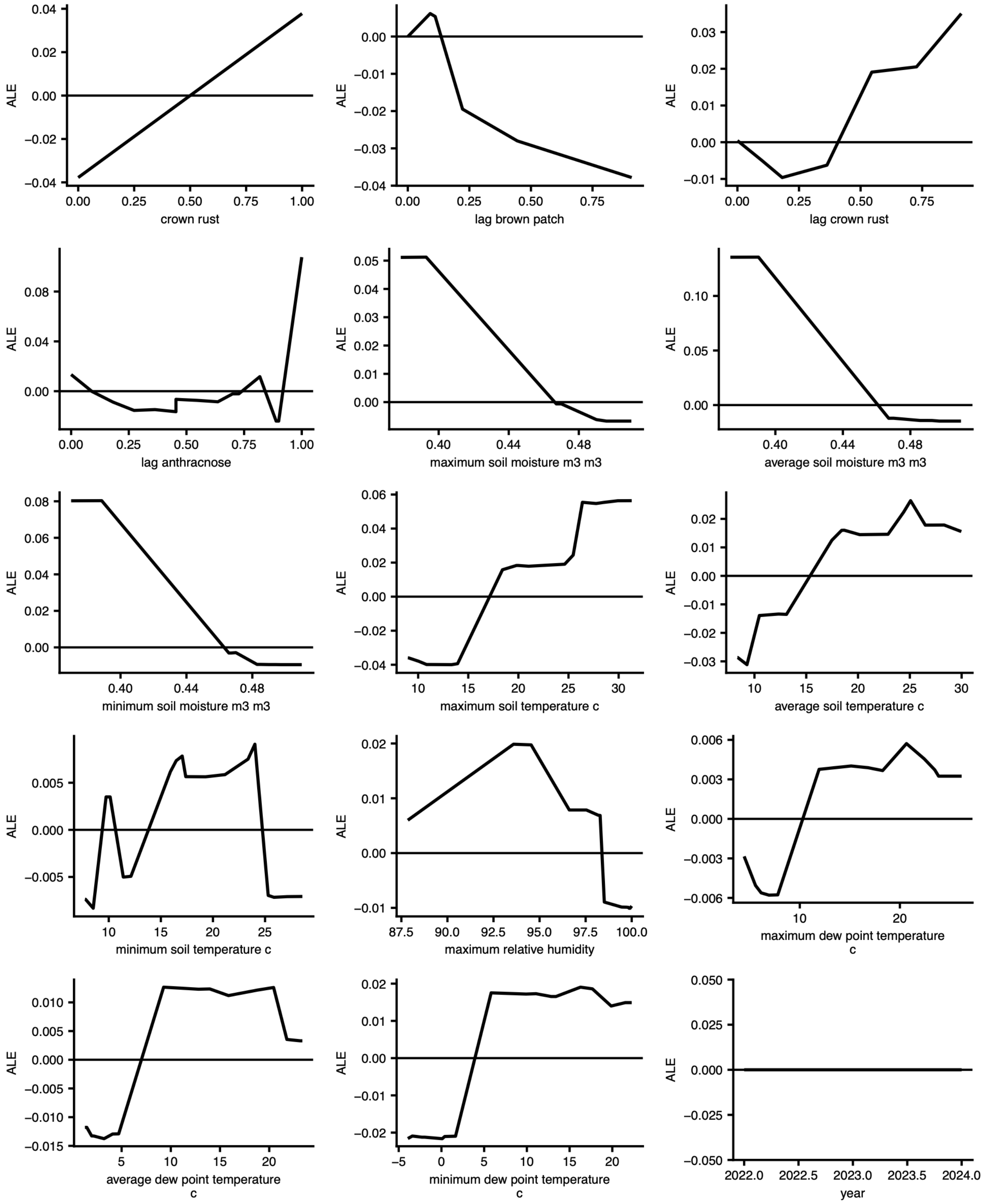

Figure S13. ALE plots for top 15 most important variables in anthracnose random forest model.

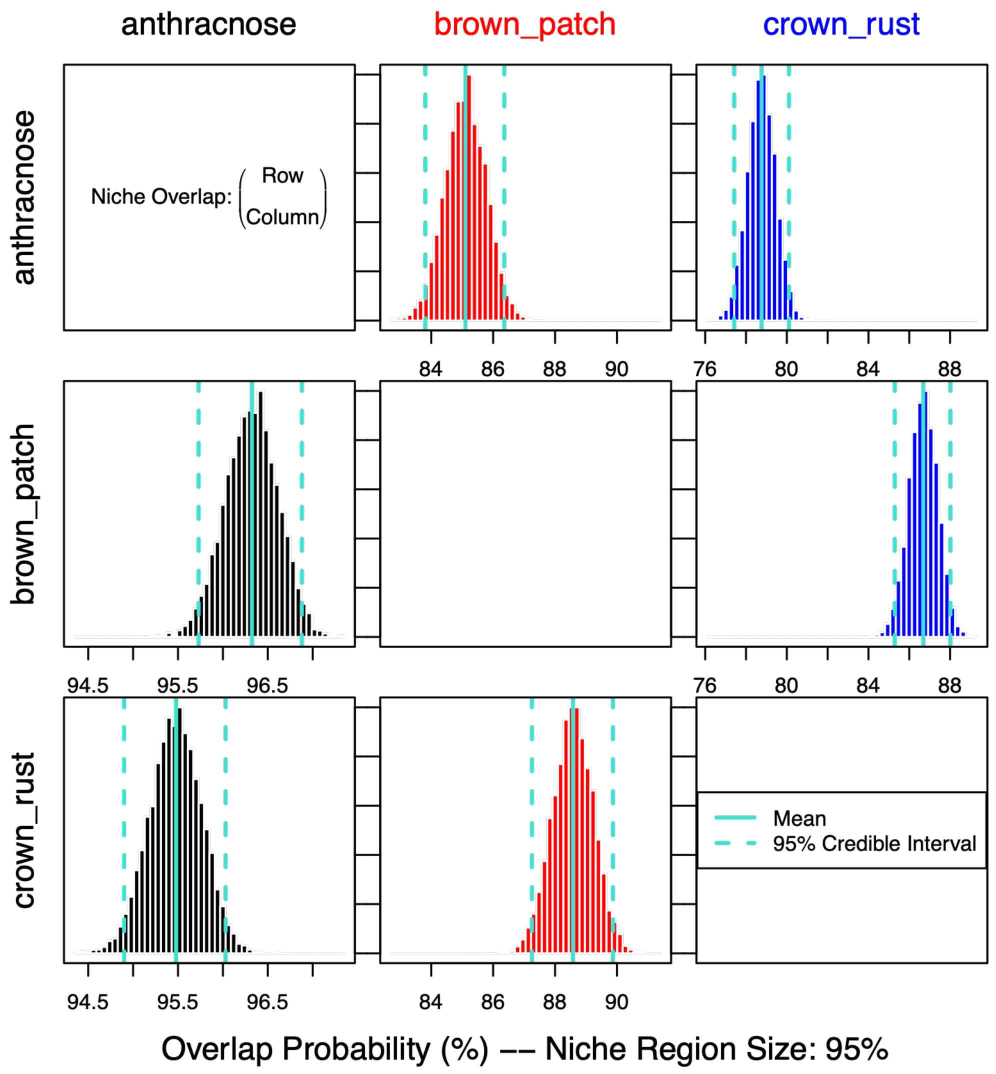

Figure S14. Niche overlap among anthracnose, brown patch, and crown rust, calculated from environmental principal components. Rows represent the focal pathogen, and columns indicate the probability that the focal pathogen occurs within the core 95% of another pathogen's environmental niche. For example, there is an 85% chance that anthracnose occurs within 95% of the environmental niche of brown patch.

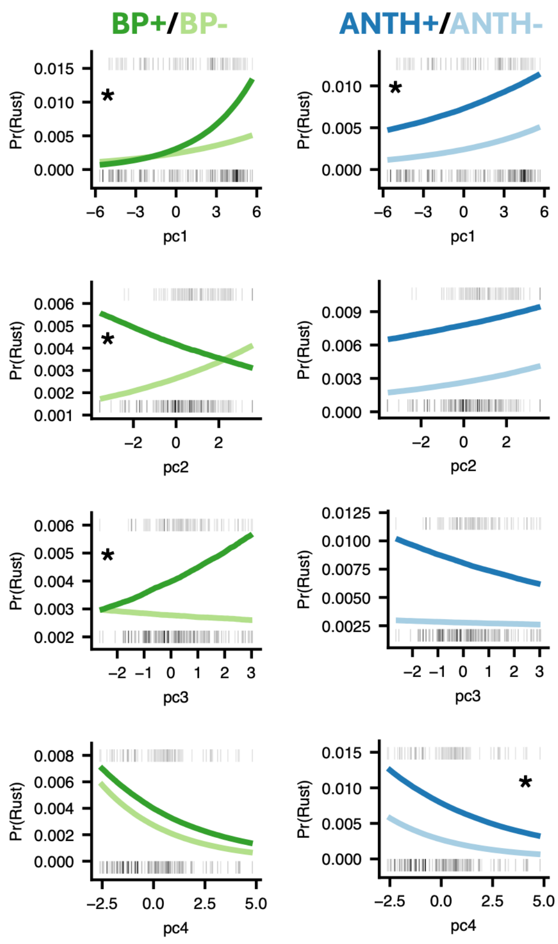

Figure S15. Conditional effects of environmental PCs and coinfection status on the probability of observing crown rust infection from Bayesian hierarchical models. Rugs along the top and bottom of each plot give the observed presences (top) and absences (bottom) of crown rust in the data across PC scores. Plots marked with (*) indicate interaction terms for which zero was not included in the 95% credibility interval of the posterior distribution. (Left) Environmental PC x brown patch conditional effect plots, where the darker green indicates brown patch coinfection. (Right) Environmental PC x anthracnose conditional effect plots, where the darker blue indicates anthracnose coinfection.

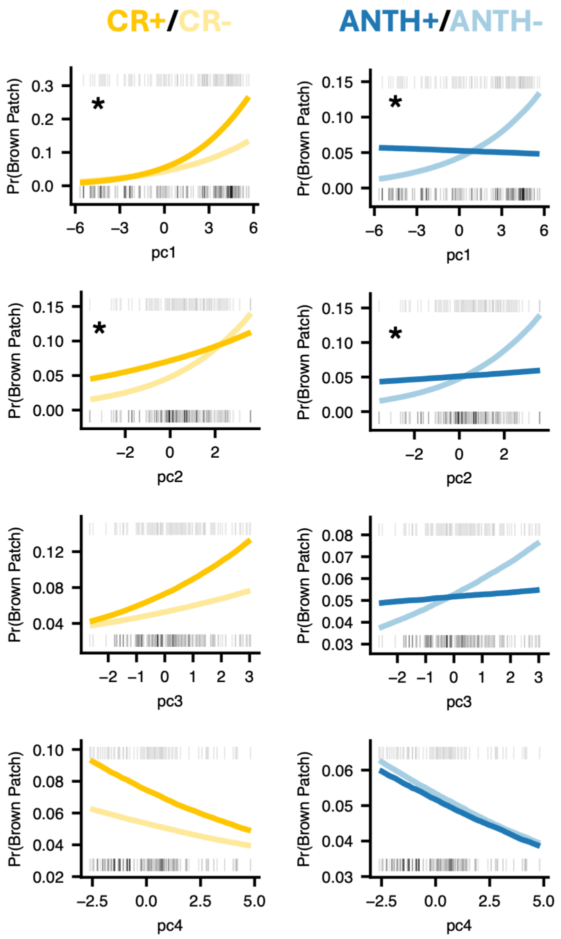

Figure S16. Conditional effects of environmental PCs and coinfection status on the probability of observing brown patch infection from Bayesian hierarchical models. Rugs along the top and bottom of each plot give the observed presences (top) and absences (bottom) of brown patch in the data across PC scores. Plots marked with (*) indicate interaction terms for which zero was not included in the 95% credibility interval of the posterior distribution. (Left) Environmental PC x crown rust conditional effect plots, where the darker yellow indicates crown rust coinfection. (Right) Environmental PC x anthracnose conditional effect plots, where the darker blue indicates anthracnose coinfection.

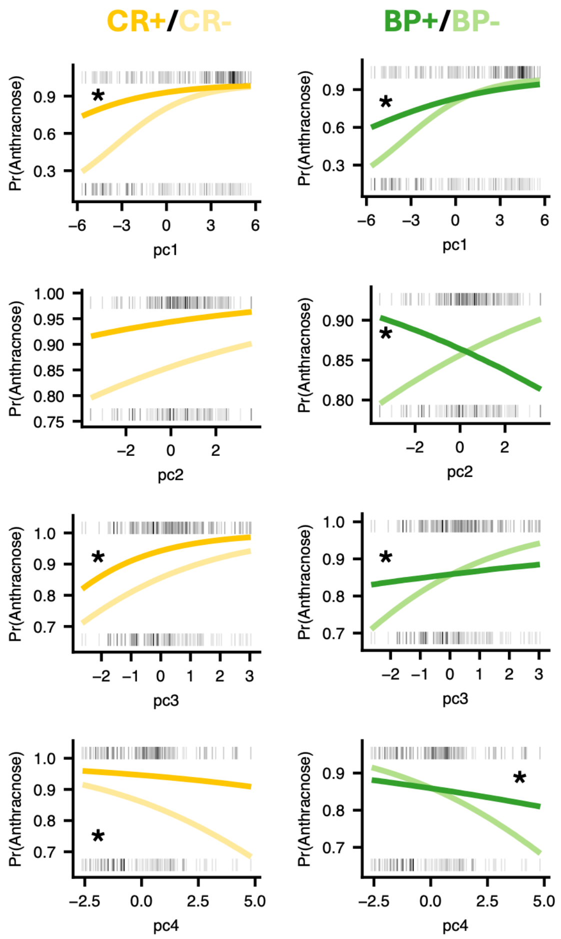

Figure S17. Conditional effects of environmental PCs and coinfection status on the probability of observing anthracnose infection from Bayesian hierarchical models. Rugs along the top and bottom of each plot give the observed presences (top) and absences (bottom) of anthracnose in the data across PC scores. Plots marked with (*) indicate interaction terms for which zero was not included in the 95% credibility interval of the posterior distribution. (Left) Environmental PC x crown rust conditional effect plots, where the darker yellow indicates crown rust coinfection. (Right) Environmental PC x brown patch conditional effect plots, where the darker green indicates brown patch coinfection.

**Supplemental Tables**

Table S1. Description of datasets included in this study.

| Dates Surveyed | Survey Level | Longitud-inal Level | Experimental Design | N Plant-Level Observations | Dataset Source |
|---|---|---|---|---|---|
| 05/08/2021 – 11/07/2024 | Plant-level | Plot-level | Monthly survey of 36 GPS-tagged locations at Widener Field. At each location, a point in a 400 $m^2$ area was located by tossing a pin flag, and then the 10 tillers nearest to the point were surveyed. | 15,371 | This Study |
| 04/15/2024 – 10/03/2024 | Plant-level | Plot-level | Twice-weekly survey of 180 1m x 1m plots. 36 plots were surveyed at a time over the course of five repetitions over the growing season. Nine tillers were surveyed per plot. Some plots were treated with fungicide and/or inoculated with Rhizoctonia during the experiment. Only the 45 plots which received no treatment were retained. | 2,432 | This Study |
| 05/24/2017 – 01/03/2020 | Leaf-level; combined to plant-level for this study | Plot-level | Monthly prevalence survey for 64 plots which had been treated year-round, for 9-months, for 7-months, or never with fungicide. Only the 16 control (never-sprayed) plots were retained. 20 random tillers were surveyed in each plot. | 9,919 | Grunberg et al., 2025: monthly_disease_survey_171819.csv |

| | | | | | |
|---|---|---|---|---|---|
| 08/15/2019 – 11/14/2019 | Plant-level | Plot-level | Halfway between the above monthly prevalence surveys, an additional 10 tillers were surveyed per plot. | 480 | Grunberg et al., 2025: midmonthly_disease_survey_2019.csv |
| 05/31/2018 – 10/31/2018 | Leaf-level; combined to plant-level for this study | Plant-level | In each plot of the fungicide treatment plots described above, 20 tillers (or the number available; as few as 5) were tagged at the beginning of the season. Tillers were then tracked weekly for disease prevalence. Only tillers from “never-sprayed” plots are included in the combined analysis (16 plots); tillers from “7-month”, “9-mont”, and “never-sprayed” treatments are included in the survival analyses (48 plots). | 6,222 (control); 13,410 (all) | Grunberg et al, 2025: longitudinal_disease_survey_2018.csv |

Table S2. Variables included in random forest models from DURH weather station.

| **Variables Retrieved from DURH** | **Unit** |
|---|---|
| Average Wind Direction | degree |
| Minimum wind speed | kph |
| Average Wind Speed | kph |
| Maximum Wind Speed | kph |
| Average Relative Humidity | % |
| Minimum Relative Humidity | % |
| Maximum Experimental Leaf Wetness | mV |
| Total Precipitation | in |
| Maximum Relative Humidity | % |
| Minimum Air Temperature | C |
| Minimum Dew Point Temperature | C |
| Maximum Dew Point Temperature | C |
| Average Dew Point Temperature | C |
| Maximum Soil Temperature | C |
| Minimum Soil Temperature | C |
| Growing Degree Days per Day - Base 50F | Degree Days |
| Average Air Temperature | C |
| Maximum Heat Index | C |
| Maximum Air Temperature | C |
| Maximum Soil Moisture | m3/m3 |
| Average Soil Moisture | m3/m3 |
| Minimum Soil Moisture | m3/m3 |
| Average Solar Radiation | W/m2 |
| Average Photosynthetically Active Radiation | W/m2 |
| Maximum Solar Radiation | W/m2 |
| Estimated Evapotranspiration (Penman-Monteith) | cm |
| Maximum Photosynthetically Active Radiation | micromol/m2s |
| Average Wind Vector - v Component | kph |
| Average Wind Vector - u Component | kph |
| Accumulated heating Degree Day (Base 65F) | degree days |

Table S3. Quantification of number of disease presence/absence observations included in random forest and Bayesian models.

| | **Absent** | **Present** |
|---|---|---|
| **Anthracnose** | 10,229 | 19,736 |
| **Crown Rust** | 24,386 | 5,559 |
| **Brown Patch** | 26,953 | 3,012 |

Table S4. (a) Model performance parameters for crown rust, anthracnose, and brown patch models with 7-day average of environmental variables leading up to sampling date. (b) Descriptions of model metrics used to assess model performance.

(a)

| ***Metric*** | ***Anthracnose*** | ***Crown Rust*** | ***Brown Patch*** |
|---|---|---|---|
| ***Accuracy*** | 0.553 | 0.853 | 0.905 |
| ***Precision*** | 0.529 | 0.952 | 0.955 |
| ***Recall*** | 0.934 | 0.878 | 0.943 |
| ***Kappa*** | 0.109 | 0.437 | 0.330 |
| ***F-Score*** | 0.676 | 0.913 | 0.949 |

(b)

| ***Metric*** | ***Description*** |
|---|---|
| ***Accuracy*** | The proportion of correct predictions made by the model. A value of 1 means the model always predicts correctly, while 0 means it always predicts incorrectly. |
| ***Precision*** | The accuracy of positive predictions, where high precision, approaching 1, indicates few false positives, while low precision, near zero, indicates many false positives. |
| ***Recall*** | How often the model predicts false negatives, where high recall (1) indicates the model did not classify any false negatives, while low recall indicates many false negatives (0). |
| ***Kappa*** | Measures how well the model predicts outcomes compared to random chance. A value of 0 indicates performance no better than chance, and 1 indicates perfect prediction. |
| ***F-Score*** | The harmonic mean of precision (the ability to avoid false positives) and recall (the ability to avoid false negatives). Higher F-Scores, closer to 1, indicate a balance of high precision and recall. This metric is particularly useful when the number of true positive cases is low, as is the case for many of our pathogens. |

Table S5. Random forest model metrics using the 14-day average of environmental variables leading up to the sampling date.

| **Pathogen** | **Accuracy** | **Precision** | **Recall** | **Kappa** | **F-Score** |
|---|---|---|---|---|---|
| Anthracnose | 0.535 | 0.524 | 0.888 | 0.0597 | 0.659 |
| Crown Rust | 0.855 | 0.912 | 0.921 | 0.393 | 0.916 |
| Brown Patch | 0.809 | 0.964 | 0.824 | 0.260 | 0.888 |

Table S6. Random forest model metrics using the 30-day average of environmental variables leading up to the sampling date.

| **Pathogen** | **Accuracy** | **Precision** | **Recall** | **Kappa** | **F-Score** |
|---|---|---|---|---|---|
| Anthracnose | 0.545 | 0.532 | 0.856 | 0.0817 | 0.656 |
| Crown Rust | 0.810 | 0.902 | 0.873 | 0.278 | 0.887 |
| Brown Patch | 0.792 | 0.965 | 0.804 | 0.242 | 0.877 |

Table S7. Full Bayesian hierarchical model results for crown rust.

| | Estimate | Est.Error | l-95% CI | u-95% CI | Rhat | Bulk_ESS | Tail_ESS |
|---|---|---|---|---|---|---|---|
| Intercept | 687 | 50 | 589 | 786 | 1 | 2820 | 2710 |
| lag_crown_rust | 2.48 | 0.0914 | 2.3 | 2.65 | 1 | 3050 | 2860 |
| month2 | -0.572 | 0.989 | -2.72 | 1.15 | 1 | 2400 | 2750 |
| month3 | -1.92 | 1.3 | -4.66 | 0.345 | 1 | 2800 | 2830 |
| month4 | -1.5 | 0.918 | -3.48 | 0.129 | 1 | 2960 | 2680 |
| month5 | -1.8 | 0.867 | -3.61 | -0.238 | 1 | 2740 | 2660 |
| month6 | 0.209 | 0.462 | -0.668 | 1.12 | 1 | 2320 | 2690 |
| month7 | 2.32 | 0.447 | 1.47 | 3.22 | 1 | 2350 | 2500 |
| month8 | 2.79 | 0.438 | 1.98 | 3.68 | 1 | 2340 | 2560 |
| month9 | 4.43 | 0.427 | 3.63 | 5.3 | 1 | 2390 | 2560 |
| month10 | 4.75 | 0.415 | 3.98 | 5.59 | 1 | 2380 | 2580 |
| month11 | 4.15 | 0.409 | 3.4 | 4.99 | 1 | 2310 | 2450 |
| month12 | 2.17 | 0.431 | 1.37 | 3.07 | 1 | 2400 | 2590 |
| year | -0.343 | 0.0247 | -0.392 | -0.295 | 1 | 2830 | 2710 |
| pc1 | 0.132 | 0.031 | 0.0717 | 0.193 | 1 | 2700 | 2790 |
| pc2 | 0.123 | 0.0522 | 0.0213 | 0.225 | 1 | 2880 | 3010 |
| pc3 | -0.024 | 0.0633 | -0.149 | 0.0987 | 1 | 2670 | 2860 |
| pc4 | -0.302 | 0.0453 | -0.393 | -0.214 | 1 | 2530 | 3000 |
| anthracnose1 | 1.15 | 0.0882 | 0.976 | 1.33 | 1 | 2980 | 3040 |
| brown_patch1 | 0.299 | 0.166 | -0.05 | 0.619 | 0.999 | 2990 | 2670 |
| anthracnose1:brown_patch1 | -0.665 | 0.138 | -0.928 | -0.393 | 1 | 3040 | 2810 |
| pc1:anthracnose1 | -0.051 | 0.0191 | -0.0884 | -0.0131 | 1 | 2900 | 2990 |
| pc2:anthracnose1 | -0.0704 | 0.051 | -0.168 | 0.0329 | 1 | 2940 | 2950 |
| pc3:anthracnose1 | -0.0645 | 0.0683 | -0.201 | 0.0744 | 1 | 2640 | 2580 |
| pc4:anthracnose1 | 0.114 | 0.045 | 0.0239 | 0.2 | 1 | 2950 | 2870 |
| pc1:brown_patch1 | 0.131 | 0.025 | 0.0812 | 0.181 | 1 | 2720 | 2810 |
| pc2:brown_patch1 | -0.205 | 0.0566 | -0.315 | -0.0936 | 1 | 2990 | 3030 |
| pc3:brown_patch1 | 0.14 | 0.0747 | -0.00655 | 0.283 | 1 | 3030 | 3080 |
| pc4:brown_patch1 | 0.0757 | 0.0516 | -0.0257 | 0.181 | 1 | 2850 | 2740 |

Table S8. Full Bayesian hierarchical model results for brown patch.

| | Estimate | Est.Error | l-95% CI | u-95% CI | Rhat | Bulk_ESS | Tail_ESS |
|---|---|---|---|---|---|---|---|
| Intercept | 1070 | 83.1 | 916 | 1240 | 1 | 2520 | 2830 |
| lag_brown_patch | 1.15 | 0.124 | 0.893 | 1.39 | 1 | 2920 | 2610 |
| month2 | 0.333 | 0.393 | -0.462 | 1.09 | 1 | 2460 | 2580 |
| month3 | 0.145 | 0.398 | -0.644 | 0.916 | 1 | 2610 | 2830 |
| month4 | 0.412 | 0.345 | -0.259 | 1.09 | 1 | 2130 | 2480 |
| month5 | 0.382 | 0.386 | -0.383 | 1.15 | 1 | 2070 | 2030 |
| month6 | 0.338 | 0.381 | -0.402 | 1.1 | 1 | 2090 | 2460 |
| month7 | 1.4 | 0.385 | 0.636 | 2.14 | 1 | 2020 | 2350 |
| month8 | 2.22 | 0.37 | 1.5 | 2.97 | 1 | 1970 | 2290 |
| month9 | 1.96 | 0.355 | 1.26 | 2.69 | 1 | 1930 | 2290 |
| month10 | 1.97 | 0.322 | 1.36 | 2.64 | 1 | 1890 | 2430 |
| month11 | 0.705 | 0.294 | 0.142 | 1.3 | 1 | 1910 | 2700 |
| month12 | 0.194 | 0.336 | -0.482 | 0.866 | 1 | 2200 | 2350 |
| year | -0.533 | 0.0411 | -0.614 | -0.455 | 1 | 2520 | 2830 |
| pc1 | 0.219 | 0.0354 | 0.149 | 0.287 | 0.999 | 2650 | 2750 |
| pc2 | 0.327 | 0.0519 | 0.225 | 0.432 | 1 | 2860 | 2950 |
| pc3 | 0.135 | 0.0678 | 0.00399 | 0.261 | 1 | 2830 | 2830 |
| pc4 | -0.0658 | 0.0476 | -0.16 | 0.028 | 1 | 2830 | 3020 |
| crown_rust1 | 0.303 | 0.152 | 0.0122 | 0.603 | 1 | 2820 | 2790 |
| anthracnose1 | 0.328 | 0.107 | 0.122 | 0.539 | 1 | 2490 | 2580 |
| crown_rust1:anthracnose1 | -0.693 | 0.137 | -0.967 | -0.435 | 1 | 2940 | 2730 |
| pc1:crown_rust1 | 0.107 | 0.0246 | 0.0594 | 0.156 | 1 | 2690 | 2740 |
| pc2:crown_rust1 | -0.19 | 0.052 | -0.292 | -0.0901 | 1 | 2890 | 2840 |
| pc3:crown_rust1 | 0.0859 | 0.0677 | -0.0505 | 0.213 | 1 | 2860 | 2810 |
| pc4:crown_rust1 | -0.0264 | 0.0484 | -0.122 | 0.0678 | 1 | 2940 | 2620 |
| pc1:anthracnose1 | -0.233 | 0.0232 | -0.28 | -0.189 | 1 | 2810 | 2790 |
| pc2:anthracnose1 | -0.28 | 0.0512 | -0.382 | -0.184 | 1 | 3040 | 2890 |
| pc3:anthracnose1 | -0.112 | 0.0715 | -0.252 | 0.0284 | 1 | 2810 | 2940 |
| pc4:anthracnose1 | 0.00237 | 0.0491 | -0.0911 | 0.101 | 1 | 2670 | 3080 |

Table S9. Full Bayesian hierarchical model results for anthracnose.

| | Estimate | Est.Error | l-95% CI | u-95% CI | Rhat | Bulk_ESS | Tail_ESS |
|---|---|---|---|---|---|---|---|
| Intercept | 232 | 42.5 | 151 | 315 | 1 | 2480 | 2460 |
| lag_anthracnose | 1.2 | 0.0781 | 1.05 | 1.35 | 1 | 2760 | 2870 |
| month2 | -0.0234 | 0.121 | -0.262 | 0.208 | 1 | 2810 | 2800 |
| month3 | -1.58 | 0.121 | -1.81 | -1.35 | 1 | 2240 | 2760 |
| month4 | -0.0903 | 0.133 | -0.358 | 0.172 | 1 | 2050 | 2590 |
| month5 | -0.942 | 0.192 | -1.32 | -0.57 | 1 | 1950 | 2650 |
| month6 | -0.184 | 0.222 | -0.603 | 0.255 | 1 | 2020 | 2770 |
| month7 | -1.26 | 0.214 | -1.68 | -0.839 | 1 | 2160 | 2480 |
| month8 | -1.55 | 0.224 | -1.98 | -1.1 | 1 | 2010 | 2290 |
| month9 | -1.77 | 0.194 | -2.15 | -1.39 | 1 | 1950 | 2560 |
| month10 | -0.91 | 0.156 | -1.2 | -0.598 | 1 | 2160 | 2620 |
| month11 | -0.585 | 0.109 | -0.799 | -0.369 | 1 | 2290 | 2750 |
| month12 | -0.85 | 0.0977 | -1.04 | -0.661 | 1 | 2790 | 2850 |
| year | -0.115 | 0.021 | -0.156 | -0.0748 | 1 | 2480 | 2460 |
| pc1 | 0.403 | 0.0209 | 0.361 | 0.445 | 1 | 2170 | 2150 |
| pc2 | 0.12 | 0.021 | 0.0794 | 0.162 | 1 | 3040 | 2690 |
| pc3 | 0.334 | 0.0273 | 0.279 | 0.388 | 1 | 2530 | 2790 |
| pc4 | -0.216 | 0.0277 | -0.27 | -0.163 | 1 | 2610 | 2870 |
| crown_rust1 | 1.16 | 0.0853 | 0.993 | 1.33 | 1 | 2990 | 3040 |
| brown_patch1 | 0.317 | 0.109 | 0.111 | 0.536 | 1 | 2930 | 2800 |
| crown_rust1:brown_patch1 | -0.71 | 0.134 | -0.965 | -0.439 | 1 | 2850 | 2670 |
| pc1:crown_rust1 | -0.132 | 0.0202 | -0.172 | -0.0914 | 1 | 3040 | 3100 |
| pc2:crown_rust1 | 0.00275 | 0.0499 | -0.0902 | 0.104 | 1 | 3070 | 2820 |
| pc3:crown_rust1 | 0.151 | 0.0599 | 0.0365 | 0.271 | 1 | 2890 | 2770 |
| pc4:crown_rust1 | 0.0987 | 0.0422 | 0.014 | 0.18 | 1 | 2890 | 2690 |
| pc1:brown_patch1 | -0.194 | 0.0243 | -0.241 | -0.146 | 1 | 2710 | 2520 |
| pc2:brown_patch1 | -0.227 | 0.0513 | -0.327 | -0.127 | 1 | 2310 | 2830 |
| pc3:brown_patch1 | -0.254 | 0.0629 | -0.377 | -0.131 | 1 | 2900 | 2940 |
| pc4:brown_patch1 | 0.143 | 0.0466 | 0.0519 | 0.236 | 1 | 2840 | 2630 |
